# Effects of Instructional Context on Neural Features of Attention during Learning Activities in Children with and without ADHD

**DOI:** 10.64898/2026.08.29.748025

**Authors:** Fang Yu Chang, Xinyi Z. Mao, Maya Khalil, Mark D. Rapport, Andrea Dillon, Sandra K. Loo, Jennie K. Grammer, Agatha Lenartowicz

**Affiliations:** Semel Institute for Neuroscience and Human Behavior, University of California, Los Angeles, United States; School of Education and Information Studies, University of California, Los Angeles, United States; Department of Psychology, University of Central Florida, Orlando, United States; Department of Psychiatry and Biobehavioral Sciences, University of California, Los Angeles, United States

**Keywords:** ADHD, attention, instructional context, aperiodic, neural oscillations, 1/f, spectral slope

## Abstract

Attention is foundational to learning, yet the extent to which features of the instructional environment differentially shape attentional engagement is not well understood. Here we leveraged mobile EEG and video-coded behavioral observation to examine attentional engagement in 6–10-year-old children with and without a diagnosis of ADHD across instructional conditions varying in delivery modality (video watching, online, in-person) and management of learning (teacher-led versus student-led). EEG measures, including alpha-band (8-12Hz) oscillations, spectral slope and offset, and behavioral measures of active engagement (AE%), passive engagement (PE%), fidgeting, off-task behavior, were examined as a function of instructional context. Attentional engagement, as indicated by higher AE%, lower alpha power, flatter spectral slope and lower offset, was greatest in student-led learning, followed by teacher-led in person learning, synchronous online learning, and asynchronous learning, respectively. Child by instructional context interactions revealed that patterns of attentional engagement were largely consistent regardless of diagnosis, with group effects observed for motor behaviors (PE%, fidgeting) but not for measures of visual attention. Similarly, age showed only main effects, whereby older children showed higher passive engagement, lower fidgeting and off-task behavior, flatter spectral slope, and lower offset. The results underscore: (i) the importance of considering environmental context, such as instructional modality and management of learning, when examining mechanisms of attention and individual differences therein, and (ii) the value of recording multiple behavioral and neural measures, as not all indices commonly assumed to reflect attention capture the same underlying processes.

## Introduction

Student attention is foundational to learning in school contexts. This is particularly evident in individuals with attention deficit/hyperactivity disorder (ADHD), which is characterized by inattention, hyperactivity, and impulsivity (American Psychiatric Association, 2022). Across development, ADHD has been associated with poorer academic achievement (Jangmo et al., 2021; Gordon & Fabiano, 2019; Kuriyan et al., 2013; Murphy & Barkley, 1996). Notably, these academic difficulties persist despite well-established ADHD treatments, which show limited success in improving school outcomes (Barkley, 1991; DuPaul et al., 2018; Rapport et al., 2000; Spooner & Pachana, 2006). One possible explanation for this discrepancy is that commonly used laboratory tasks of attention do not fully capture the attentional demands encountered in everyday learning environments. Indeed, laboratory assessments of inattention show weak associations with classroom observations and real-world academic functioning (Barkley, 1991; McGee et al., 2000). Collectively, these findings suggest that attention in naturalistic settings is shaped by contextual factors not present in laboratory settings.

In a natural environment like a classroom, attention is subject to multiple factors – such as sensory distraction (e.g., talking), level of arousal (e.g., fatigue), and motivation (e.g., boredom) – and variable attentional support from instructional scaffolding or self-regulation. These environmental dynamics impact behavior and academic performance as illustrated in a series of studies by Connor and colleagues (Connor et al., 2009; Connor & Morrison, 2016; Morrison et al., 2005), which revealed significant differences in learning outcomes as a function of: (a) instructional management (teacher-managed vs. student-managed), (b) group structure (whole class, small group, individual), (c) content, and (d) learning duration. Teacher-managed, small-group instruction was particularly noteworthy and reported to be nearly 4 times more effective in improving literacy skills in preschool than the same type of instruction provided to an entire class (Connor et al., 2006, 2011, 2013). These findings align with ADHD research (Imeraj et al., 2013, 2016), whereby small-group activities improved on-task performance in children with ADHD (relative to whole-group or individual contexts) independent of teacher-supervision, and show susceptibility to duration of “idle” time during classroom activities (Imeraj et al., 2016). In complement, duration of instruction is likely to play a significant role in learning outcomes based on evidence that in-class on-task performance in 6-11 yr olds differ significantly between children with (2-4 min) and without ADHD (7 min) (Rapport et al., 2009), both of which are lower than the 10 min duration suggested by others.

Consequently, a key factor to understanding the mechanisms and processes that improve student attention, including in ADHD, is assessment of attention across learning contexts using objective measures. Mobile electroencephalography (EEG) has emerged as a desirable tool for quantifying children’s attention (Niso et al., 2023), with increasing application in the educational domain (García-Monge et al., 2024). Of particular importance is the availability of a reliable EEG indicator of visual attention, namely alpha-band oscillations (8-12 Hz). At rest, increases in alpha power (i.e., squared magnitude of the oscillations) are thought to indicate disengagement of visual attention (Pfurtscheller, 1992; Sadaghiani & Kleinschmidt, 2016), while event-related decreases of alpha power are coupled with visual encoding (Foxe & Snyder, 2011; Klimesch, 2012; Mathewson et al., 2011; McWhirter & Klapper, 1990), associated with neural substrates including occipital, thalamic and fronto-parietal interactions (de Munck et al., 2007a; Laufs et al., 2003; Lenartowicz et al., 2016; Scheeringa et al., 2009). Differences in modulation of alpha power during attention tasks have also been reported in children with ADHD (Lenartowicz et al., 2018; Michelini et al., 2022).

In the present investigation, we leveraged mobile EEG and video-coded behavior to quantify the extent to which attention engagement in children with and without ADHD was optimized while engaged in learning activities under varying instructional modalities and management contexts.

## Materials & Methods

### Participants

Eighty participants enrolled in this multi-site study, approved by IRBs of University of California, Los Angeles (19-000139) and University of Virginia (4956). Flyers were distributed to recruit participants in the greater Los Angeles and Charlottesville area. Children were eligible to participate if they were between ages of 6 and 10 years. The exclusion criteria were presence of serious medical or neurological illnesses likely to influence cognition or brain function (e.g., epilepsy, head trauma, or other neurologic disorders), and presence of clinically unstable conditions (e.g., suicidality, psychosis, mania, significant aggression, substance abuse). Both participants’ assent and parents’ consent were obtained prior to data collection. Participants were referred to a licensed clinician to receive a semi-structured assessment for ADHD if their Conners’ Inattention and Hyperactivity T-scores were over 70. The detailed protocol is provided in the supplementary materials.

### Data Collection

Following initial screening and consent, participants and their parent/caregiver were invited for a single visit to one of the two study sites. During the visit, the parent/caregiver completed a series of neurocognitive and diagnostic questionnaires, while the child completed computerized cognitive tasks and learning activities that varied in instructional modality and instructional management as described below, during which EEG and video of child behavior were recorded. This was followed by a set of academic achievement assessments. The total session was approximately 2.5 hours including session orientation, EEG cap set-up, data collection, and short breaks between tasks.

### Experimental Manipulation of Instructional Context

Following consent and orientation, participating children were invited to engage in a series of neuroscience-themed activities about the brain and neurons. Drawing on the individualized student instruction (ISI) framework (Connor et al., 2004), four learning activities, which varied in instructional modality and instructional management, were designed to mimic real-world learning experiences as detailed below.

*Instructional modality* was varied among three teacher-led activities: online video watching, online instruction, and in-person instruction. The online and in-person instruction activities varied in whether learning was conducted online or in-person. In both activities, the participant listened to content provided by an instructor and responded to questions; however, while in the in-person activity the instructor sat next to the participant at the table, in the online activity, the instructor moved to a different room and all interactions occurred online. The online instruction activity was also compared with online video watching activity to determine whether attention varies when learning is synchronous (interacting with the instructor) or asynchronous (video watching). We hypothesized that attention would increase from the video watching (online, asynchronous), to the online instruction (online, synchronous), to the in-person instruction (in-person, synchronous), given the increasing level of interaction with either the content or the teacher.

A fourth activity—individual learning—was included to test the effect of *instructional management*, relative to the in-person instruction activity. Namely, while in the in-person activity learning was teacher-led, in the individual activity, the participant (the student) led the activity by creating a neuron using arts and crafts materials (i.e., pipe cleaners, beads, and clay) with minimal technical help from the instructor. This individual activity (in-person, synchronous, *student-led*) was compared with the in-person teaching context (in-person, synchronous, *teacherle*d) to assess whether benefits to attention occurred from scaffolding provided by the teacher, which we predicted would be most beneficial to children with ADHD.

Each activity was designed to be 10-minutes long and was delivered one-on-one by a trained instructor. Due to natural variations in the pace of interaction and the time needed to complete the individual task, the duration of activities varied among participants, with an average of 7.00 minutes for the online, 6.49 minutes for the in-person, and 8.42 minutes for the individual. The video watching activity was consistently 10-minute because it was the same clip for all participants and did not involve any interaction. The four instructional contexts were presented in four different orders to control for possible order effects (see supplementary materials for more information).

### Computerized Cognitive Tasks

The participant performed 4 computerized tasks prior to the learning activities for comparative analysis (see supplemental materials for full description). Relevant to the current study, eyes-closed (EC) and eyes-open (EO) were 1-min each, with a black screen during EC and a centered snowflake during EO. These resting-state conditions served as baseline references for EEG measures during the learning activities.

### EEG & Video Recording

Throughout the entire session we recorded both EEG signals and video. The EEG signal was digitized at 250 Hz using Smarting Mobi (mBrain Train, Serbia) with a 24-channel saline cap (Greentek Pty Ltd, Wuhan, China). The impedance of each electrode was < 40 kW. Participants were encouraged to remain as still as possible during the EEG recording, but movement was not impeded or corrected. We also collected video recordings for each participant from the front and side angles, subsequently used for behavioral coding. The videos were recorded through two webcams (Logitech HD Pro Webcam C920, Logitech) with Logitech Capture software (Logitech).

### Data Processing & Feature Extraction

We used custom scripts in MATLAB (The MathWorks, 2022) and EEGLAB (Delorme & Makeig, 2004) to process the data. Preprocessing steps included 0.1Hz high-pass and 50Hz low-pass filters, artifact subspace reconstruction for non-stationary artifacts, and independent components analysis for automated removal of muscle, eye, heart, line noise and channel artifacts (full details are provided in supplementary materials). Clean time series were segmented into 2-s non-overlapping epochs. Noisy epochs were rejected using a 5-SD threshold and pop_autorej.m (≤3% rejection or at least one epoch). Dataset exclusion criteria included: RMS outside the IQR, >50% rejected epochs, or <10 remaining epochs. Fifty-nine participants were retained after exclusions due to age, non-participation, missing diagnostic data, excessive noise, data loss, equipment failure, insufficient trials, or RMS outliers. On average, retained epochs were 23 (EC), 22 (EO), 242 (video), 187 (online), 169 (in-person), and 213 (individual), with an average retention rate of 82%.

The primary feature of interest— power of alpha-band oscillation (8-12Hz)—was hypothesized to decrease with increased visual engagement. To extract alpha power, the power spectral density (PSD) for each frequency was calculated for each epoch using pwelch function of MATLAB without overlapping. In addition, we quantified the distribution properties of the entire spectrum, calculating both the spectral slope (proportional to the slope of the log transformed spectrum) and offset. These measures were included as exploratory metrics, based on the hypothesized value of the relative distribution of oscillatory frequencies in varying with brain state. For instance, the spectral slope has been found to be flatter in adolescents with ADHD (Karalunas et al., 2022; Ostlund et al., 2021) and hypothesized to be associated with increased neural noise (Pertermann et al., 2019). As such, we hypothesized that a flatter spectral slope would be observed in children with ADHD and during less engaging conditions.

To extract power independently from spectral slope and offset, the extracted PSD was input into the spectral parameterization or specparam (formerly known as “Fitting Oscillations and One-Over-f” or FOOOF) toolbox (Donoghue et al., 2020). Finally, the extracted data (power, spectral slope & offset) were averaged across the channels. Specifically, nine channels were selected for the alpha oscillation analysis, and grouped into three distinct clusters to increase signal-to-noise ratio (SNR) while covering the anterior to posterior scalp: F3, F4, and Fz were grouped as the frontal cluster; C3, C4, and Cz were grouped as the central cluster; O1, O2, and POz were grouped as the posterior cluster. Results for frontal and central clusters are presented in supplemental materials for brevity, as are results for spectral offset, which were consistent with but more variable than the other features.

### Video-Based Behavioral Coding

We used video-based behavioral coding to quantify attention-related behaviors throughout the activities. Namely, second-by-second behavior was categorized as active or passive engagement, fidgeting, or off-task gaze/verbalizations as described in Table 1 (full details in supplemental materials). For each behavioral category/code during each activity, a percentage was generated to capture the proportion of time a child exhibited a particular behavior in relation to the total length of the activity. For example, during a 10-minute individual activity, if the participant had 7 minutes coded as actively engaged in making the neuron, the percentage of AE during the individual activity would be (7 minutes /10 minutes) * 100% = 70.00%.

**Table 1.** Video-Based Behavioral Coding Categories.

| Category | Code | Definition | Examples | Adapted From |
| --- | --- | --- | --- | --- |
| Engagement | Active Engagement | Actively engaging with the task. | Answering instructor's questions; following the task or instructor's directions. | Shapiro (2004) |
|  | Passive Engagement | Attending to the task without active interaction. | Listening to the instructor; watching a visual display without conversing; nodding to show understanding. | Shapiro (2004) |
| Fidgeting | Macro Fidgeting | Moving the head, appendages, or body with a complete spatial displacement relative to a starting position, when not required by the task. | Raising hands over the head; sliding down in the chair. | Farley et al. (2013) |
|  | Micro Fidgeting | Exhibiting repeated or small movements with minimal spatial displacement. | Shaking feet or legs; playing with fingers; tapping a pencil on the table. | Farley et al. (2013) |
| Off-Task | Passive Off-Task | Showing gaze away from the task or instructor for two or more seconds. | Looking at the door; peeking at the recording cameras; staring out the windows. | Rapport et al. (2009) |
|  | Verbal Off-Task | Making audible verbalizations unrelated to the task. | Making random noises; asking task-irrelevant questions. | Shapiro (2004) |

### Behavioral and Cognitive Measures

Caregiver-reported ADHD symptoms were assessed using the Conners (Conners et al., 2011). Attention and behavioral regulation were measured using the SWAN(Lakes et al., 2012). Academic achievement and processing speed were assessed using selected subtests of the Woodcock–Johnson IV (WJ-IV; Schrank et al., 2014a; Schrank et al., 2014b), including Letter-Word Identification, Passage Comprehension, Applied Problems, Math Facts Fluency, and Pair Cancellation.

### Group Statistics

Our primary objective was to evaluate the effects of diagnostic group and instructional context across activities on EEG features and behavioral data. To do so, we used a repeated measures ANOVA (activity * group) with age and gender as covariates. There was no interaction between activity and age or gender either in the EEG features or behavioral data. Consequently, age and gender were removed for the remaining analyses (cf., supplementary materials).

Due to inequalities in sample size between participants with ADHD (N = 15) and without ADHD (typically developing, TD; N = 44), we conducted a secondary analysis in which groups were constructed based on T-scores ≥ 60 for inattentive and hyperactivity symptoms, dividing participants into high (N = 29) and low (N = 30) symptom groups. The results of this analysis were comparable with those of original diagnostic label grouping and thus are not presented in the main text. However, as they provide results with balanced samples for statistical purposes, they are included in the supplementary materials.

### Brain-Behavior Analyses

To test the relationship of the observed effects to behavior, we incorporated a 2-step analytic procedure to assess time-on-task effects and within-subject brain-behavior relationships. For the initial step, we used post hoc linear contrasts to evaluate time-on-task effects for each outcome variable across the four quarters of each learning activity, which allowed us to assess whether video-coded behavior and EEG features of attention similarly tracked time-on-task effects on engagement (e.g., fatigue). The second step, assessed the *within-subject* correlation between alpha power and video-coded behavior by calculating a within-subject slope, regressing each video-coded outcome on alpha power across the entire session (across all four activities), and evaluating the group average of the within-subject slopes using a one-sample t-test.

Finally, we conducted several additional analyses to assess potential confounding effects. To test for inequalities in gender, a Chi-square test was utilized in the subscales of Conners, SWAN and WJ subscales. Additionally, an independent Samples T-test was used to test for inequalities between the groups in the subscales of Conners, SWAN, WJ, the length of each activity, and the epoch numbers. Across all analyses, the significance level was set at 0.05, and IBM SPSS Statistics (IBM Corp., 2022) was utilized for all statistical analyses.

For primary analyses, effects were evaluated within a repeated-measures ANOVA (activity * group), which controls the family-wise error rate at the model level. Follow-up pairwise comparisons were limited to a small number of planned contrasts within factors and were conducted using the least significant difference (LSD) approach as implemented in SPSS. Given the restricted number of comparisons and the structure of the model, additional correction for multiple comparisons was not applied. Secondary analyses (independent-samples t-tests) assessing potential confounding effects were exploratory and interpreted with caution.

### Design Considerations and Sample Size

The present study employed a cross-sectional design to examine associations between instructional context, behavioral engagement, and neural measures of oscillatory activity. The sample size was determined based on prior EEG studies (de Munck et al., 2007b; Gibbings et al., 2021), which reported large effect sizes (Cohen’s *d* ≈ 0.7–0.81), although these effects may vary when groups are defined based on inattentiveness. Moreover, effects of instructional or activity context remain less well characterized. We therefore adopted a conservative a priori small-to-medium effect size (*f*^*2*^ = 0.2) for power estimation. Given the lower bound of our sample size and alpha = 0.05, the design provides > 0.80 power to detect between-subject effects and > 0.95 power to detect within-subject effects and interactions.

## Results

### Participant Demographics & Assessments

As expected, inattention (*p* < 0.001) and hyperactivity symptoms were significantly higher in the ADHD relative to the TD (*p* < 0.001) participants on both Conners and SWAN scales (Table 2). Groups differed significantly in a subset of academic achievement scores, with scores on letter-word identification, applied problems and math facts fluency significantly lower in the ADHD than in the TD (*p* < 0.05) participants; however, no significant differences were found in passage comprehension and pair cancellation. Additionally, there was no significant difference in sex distribution or age between the ADHD and TD groups (*p* = 0.38 and *p* = 0.35 respectively).

**Table 2.** Participant Demographics.

|  | ADHD (M, SD) | TD (M, SD) | Statistic | *p* | Effect size |
| --- | --- | --- | --- | --- | --- |
| N | 15 | 44 | - | - | - |
| Sex (male : female) | 10:5 | 23:21 | $\chi^2(1) = 0.94$ | 0.33 | $V = 0.13$ |
| Age (year) | 8.24 (1.36) | 8.63 (1.37) | $t(57) = -0.95$ | 0.35 | $d = -0.29$ |
| **ADHD Rating Scale Measure** |  |  |  |  |  |
| Conners - Inattention scale | 83.87 (7.05) | 52.39 (8.77) | $t(57) = 12.57$ | < 0.001 | $d = 3.76$ |
| Conners - Hyperactivity scale | 78.40 (10.43) | 51.25 (7.78) | $t(57) = 10.67$ | < 0.001 | $d = 3.19$ |
| SWAN - Inattention scale | 1.04 (0.63) | -0.49 (0.92) | $t(57) = 5.98$ | < 0.001 | $d = 1.79$ |
| SWAN - Hyperactivity scale | 0.93 (0.91) | -0.59 (0.79) | $t(57) = 6.19$ | < 0.001 | $d = 1.85$ |
| **Academic achievement (WJ)** |  |  |  |  |  |
| Letter-Word Identification | 42.40 (16.69) | 53.98 (14.51) | $t(57) = -2.57$ | 0.013 | $d = -0.77$ |
| Passage Comprehension | 24.00 (9.08) | 28.98 (8.51) | $t(57) = -1.92$ | 0.06 | $d = -0.58$ |
| Applied Problems | 26.73 (7.06) | 31.66 (6.05) | $t(57) = -2.61$ | 0.012 | $d = -0.78$ |
| Math Facts Fluency | 32.87 (22.00) | 49.55 (24.85) | $t(57) = -2.31$ | 0.025 | $d = -0.69$ |
| Pair Cancellation | 40.53 (17.53) | 43.22 (12.89) | $t(57) = -0.64$ | 0.53 | $d = -0.19$ |

Note. *N* = number, *M* = mean, *SD* = standard deviation. $\chi^2$ tests were used to examine sex differences between groups; independent-samples *t* tests were used to examine group differences in age, ADHD symptom measures, and WJ scores. Effect sizes are reported as Cramér's *V* for $\chi^2$ tests and Cohen's *d* for *t* tests.

### Behavioral Engagement

To identify differences in effects of instructional context on attention, we examined video-coded behavioral engagement. Participants’ engagement levels were assessed independently for each coded category (Figure 1): active engagement (AE), passive engagement (PE), fidgeting and off-task behavior.

**Figure 1.**
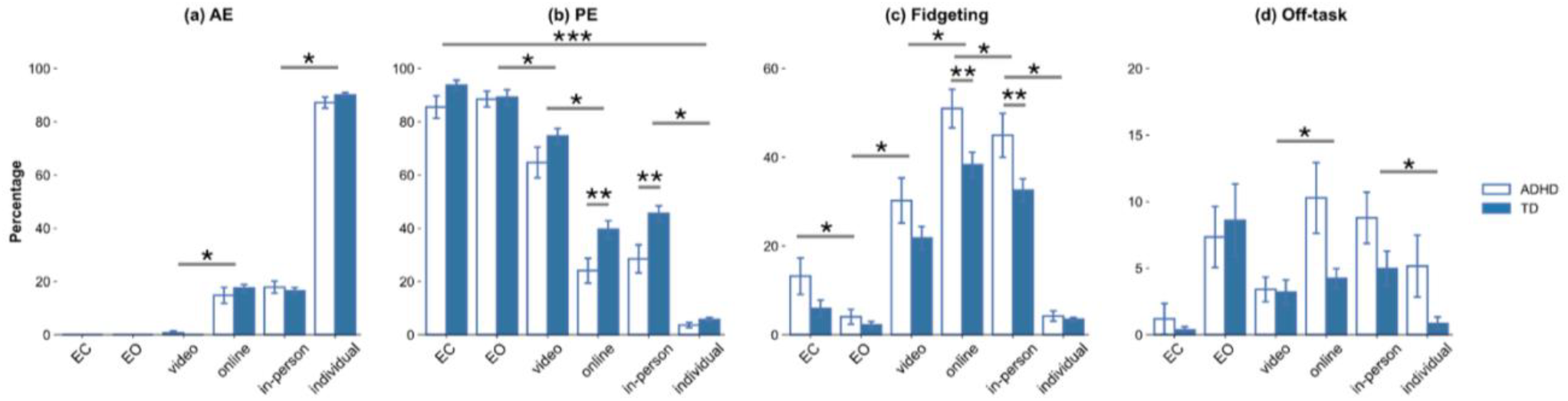
Video-coded behavior across instructional context. Main effects of activity indicate the greatest active engagement (AE%), and lowest passive engagement (PE%), fidgeting, and off-task behaviors during the student-led (individual) activity. Notably, fidgeting (and the negatively correlated PE) was greater in children with ADHD relative to TD children during the online and in-person activities. Off-task behaviors showed a similar but less reliable pattern of effects. * *p* < 0.05, main effect of activity. ** *p* < 0.05, main effect of group. *** *p* < 0.05, interaction between activity and group; TD = typically developing. Error bars represent standard error.

*Active Engagement (AE%)*. Neither the main effect of group (*F*(1,55) = 0.69, *p* = 0.41, η^2^_p_ = 0.012) nor the interaction between activity and group was significant (*F*(5, 275) = 1.24, *p* = 0.29, *η*^2^_p_ = 0.002). However, there was a significant main effect of activity (*F*(5, 275) = 42.68, *p* < 0.001, *η*^2^_p_ = 0.44). AE% increased across activities (left to right in Figure 1(a)), whereby, based on the post-hoc pairwise tests, AE% was the highest in the individual activity relative to all other activities (*p* < 0.001). AE% was also higher in the in-person learning activity relative to the video watching, EO and EC (*p* < 0.001), and there was no significant difference between AE% in the online and inperson activities (*p* > 0.05). The results suggest that attention engagement was the strongest in the student-led activity, followed by teacher-led activities (online, in-person), and finally asynchronous online learning (video watching).

*Passive Engagement (PE%)*. Consistent with AE% results, there was a main effect of activity (*F*(5,275) = 5.451, *p* < 0.001, *η*^2^_p_ = 0.09), PE% decreased across activities (left to right in Figure 2b). Post-hoc pairwise results showed that PE% was the highest in EC relative to other activities (*p* < 0.001), but not for EO (*p* > 0.05). PE% was also higher in EO relative to instruction contexts (*p* < 0.001). Within learning activities, video watching had the highest PE% among all activities (*p* < 0.05), while the individual activity had the lowest PE% (*p* < 0.001). Unlike AE%, there was a significant interaction between activity and group (*F*(5,275) = 2.37, *p* = 0.03, *η*^2^_p_ = 0.041) for PE%, as well as a significant main effect of group (*F*(1,55) = 5.97, *p* = 0.01, *η*^2^_p_ = 0.098). As shown in Figure 2(b), ADHD had lower PE% than TD in the online and in-person activities (*p* < 0.05). Thus PE% showed an overall trend opposite to AE% across activities, consistent with a negative relationship between active and passive behavior. Additionally, and unlike AE%, PE% differentiated groups across instruction modality in the teacher-led activities (online & in-person).

**Figure 2.**
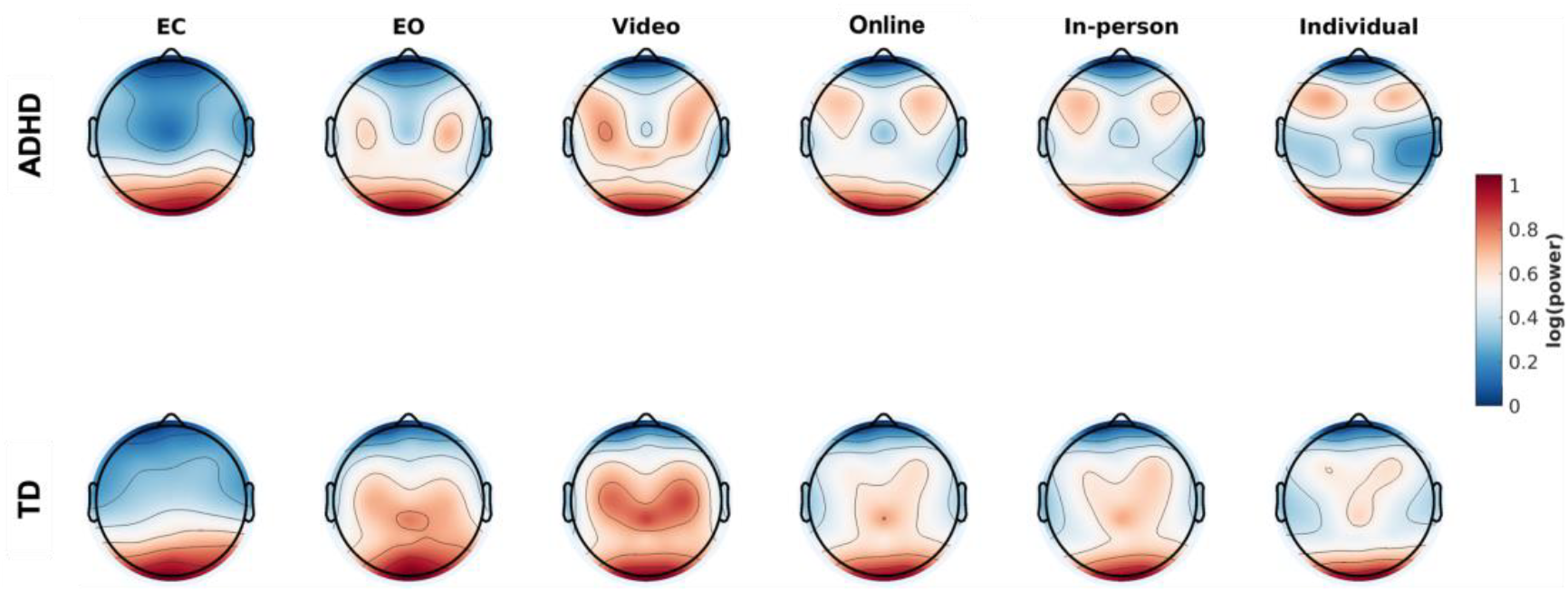
Topographical distribution of alpha power across resting state and activities divided in groups. Alpha power was elevated during the resting state (EC, EO) relative to the learning activities, with an occipital scalp distribution. During the learning activities, alpha power was observed across both central and posterior scalp regions, decreasing progressively from asynchronous (video watching) to teacher-led synchronous (online, in-person) to student-led (individual) activities. TD = typically developing.

*Fidgeting*. There was no significant interaction between activity and group (*F*(5, 275) = 1.60, *p* = 0.16, *η*^2^_p_ = 0.028) in fidgeting measures, however, there was a main effect of group (*F*(1,55) = 7.40, *p* = 0.009, *η*^2^_p_ = 0.119 and activity (*F*(5,275) = 5.74, *p* < 0.001, *η*^2^_p_ = 0.095). Fidgeting was significantly more frequent in ADHD than TD in the online and in-person activities (*p* < 0.05) (Figure 1(c)). Additionally, the post-hoc pairwise results showed that fidgeting was the highest in the online activity relative to among all activities (*p* < 0.05). Fidgeting during the In-person activity was also elevated relative to the video watching and individual activities (*p* < 0.001). There was significant difference in fidgeting between EC, EO, and the individual activities (*p* < 0.05). Collectively, fidgeting behavior shows effects analogous to, but opposite in direction, to the PE% metric, revealing the highest level of fidgeting (the lowest PE%) during the teacher-led, synchronous activities (online, in-person) and asynchronous learning (video watching) relative to the student-led (individual) context or resting state (EC, EO). Similar to the PE% results reported above, fidgeting was significantly, higher in children with ADHD relative to TD children.

*Off-Task*. For measures of off-task gaze/verbalizations, neither the main effect of group (*F*(1,55) = 1.56, *p* = 0.21, *η*^2^_p_ = 0.028) nor the interaction between activity and group was significant (*F*(5,275) = 1.70, *p* = 0.13, *η*^2^_p_ = 0.030). However, there was a significant main effect of activity (*F*(5,275) = 5.14, *p* < 0.001, *η*^2^_p_ = 0.086) (Figure 2(d)). The post-hoc pairwise results revealed that off-task behaviors in the video watching were significantly lower relative to the online (*p* < 0.001) and in-person (*p* = 0.004) learning activities. There was no significant difference between the online and in-person activities (*p* > 0.05), but off-task% was significantly higher during the In-person relative to the individual activities condition (*p* = 0.001).

There was a between-subject effect of age in passive engagement (*F*(1,55) = 9.41, *p* = 0.003, *η*^2^_p_ = 0.146), fidgeting (*F*(1,55) = 5.81, *p* = 0.02, *η*^2^_p_ = 0.096), and off-task behavior (*F*(1,55) = 8.87, *p* = 0.004, *η*^2^_p_ = 0.139). Older participants showed higher PE% and lower fidgeting and off-task behavior. No interactions were observed with instructional context manipulations.

In sum, across measures, the student-led (individual) instructional context showed the greatest behavioral indicators of engagement, followed by teacher-led contexts (online, in-person) and, finally, asynchronous learning (video watching), suggesting that instructional management and modality both impact attention engagement during learning. Notably, only passive engagement and fidgeting differentiated the groups during the teacher-led activities, whereas between-group differences were not significant in metrics of active engagement or off-task behavior.

### EEG Extracted Features During Learning Activities

Figure 2 shows overall alpha power distributions over the scalp across instructional contexts. The statistical results are reported here for the posterior cluster, whereas the results for the central and frontal clusters are listed in the supplementary material, as results were analogous in directionality.

*Alpha Power*. Across posterior electrodes, neither the main effect of group (*F*(1,55) = 1.94, *p* = 0.16, *η*^2^_p_ = 0.034), nor interaction between activity and group were significant (*F*(5,275) = 0.55, *p* = 0.73, *η*^2^_p_ = 0.01). Analogous to the AE% results, the posterior cluster showed only a significant main effect of activity (*F*(5,275) = 5.56, *p* < 0.001, *η*^2^_p_ = 0.092) as alpha power decreased across two resting state conditions and activities (from left to right in the upper panel of the Figure 3(a)). The post-hoc pairwise results indicated that alpha power was the highest in EC relative to all learning activities (*p* < 0.001), as expected with elimination of visual input to visual cortices. Alpha power was also higher in EO relative to all the learning activities (*p* < 0.001), suggesting additional visual engagement during the activities. Among instructional contexts, consistent with video-coded AE% measures, alpha power was the highest during the video watching activity (*p* < 0.001) relative to other activities, and the lowest during the individual activity (*p* < 0.001) relative to other activities. However, the post-hoc pairwise results showed that alpha power between the online and in-person was not significantly different (*p* > 0.05). In sum, the stepwise decrease in power among the learning activities suggests a progressive increase in visual engagement from asynchronous (video watching) to teacher-led (online & in-person) to student-led (individual) contexts, which is consistent with the effects on video-coded behavior and implies that both instructional modality and instructional management affect attention during learning.

**Figure 3.**
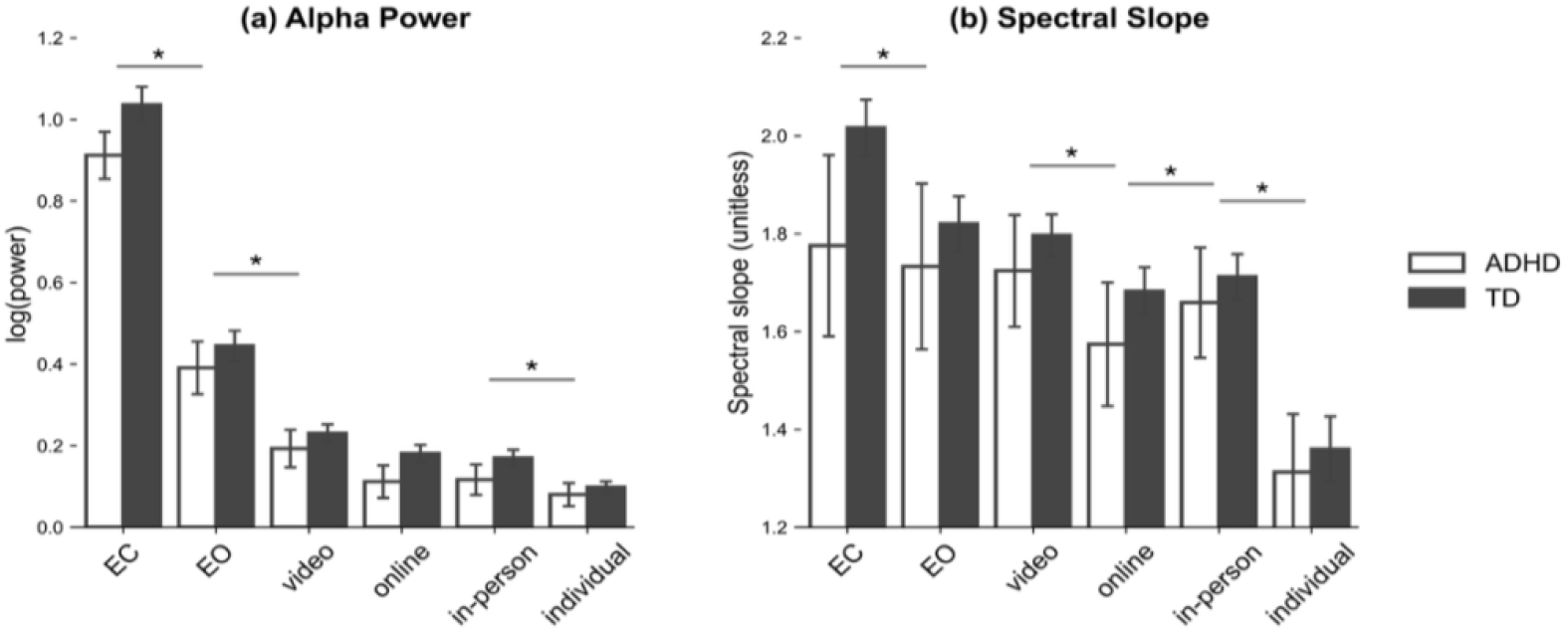
Average (a) alpha power, (b) spectral slope in posterior (O1,O2, POz) electrode clusters. Across analyses, main effects of group were not significant. Alpha power (Figure 3(a)) decreased from the resting state (EC > EO), to asynchronous (EO > video watching), to teacher-led synchronous (video watching > online/in-person), and finally to the student-led (individual) activities. A similar pattern was observed for slope (Figure 3(b)), suggesting a flattening of the power spectrum with alpha power decrease. * *p* < 0.05, main effect of activity. ** *p* < 0.05, main effect of group. *** *p* < 0.05, interaction between activity and group. TD = typically developing. Error bars represent standard error.

*Spectral Slope*. The spectral slope, which represents the log slope of the spectral density function, showed results consistent with those of alpha power. Neither the main effect of group (*F*(1,55) = 1.56, *p* = 0.21, *η*^2^_p_ = 0.028) nor the interaction between activity and group were significant (*F*(5,275) = 1.55, *p* = 0.18, *η*^2^_p_ = 0.027). As for alpha power, the posterior cluster showed only a significant main effect of activity, with the slope decreasing across least to most engaging activities (*F*(5,275) = 2.94, *p* = 0.01, *η*^2^_p_ = 0.051) (from left to right in the upper panel of Figure 3(b)), which indicates that the spectral slope is flattening (relative decrease in low frequency power, increase in high frequency power). The post-hoc pairwise results indicated that the slope was the highest (slope was steepest) in EC relative to all other activities (*p* < 0.05). The slope was also higher in EO relative to all learning activities (*p* < 0.05), except for video watching and in-person (*p* > 0.05). Among activities, the slope was the highest (steepest) during the video watching activity (*p* < 0.001) relative to other activities and lowest during the individual activity (*p* < 0.001) relative to other activities. In sum, the spectral slope largely tracks effects in alpha power across activities, and additionally indicates a flattening of the spectral slope, from least to the most engaging activities. Note that the similarity in effects across measures is consistent with significant correlations among them across individuals and three clusters (ρ = .26–.92, all *p*s < 0.05).

There was a between-subject effect of age on posterior spectral slope (*F*(1,55) = 5.30, *p* = 0.03, *η*^2^_p_ = 0.088). Older participants showed flatter spectral slope. There were no significant interactions with manipulations of instructional context.

### Brain-Behavior Correlations

To test for a relationship between the EEG metrics and behavioral indicators we assessed time-on-task effects and within-subject correlations between brain and behavior measures. These are described in turn. Note that effects are presented for the entire sample to maximize power given limited sample size.

### Task-on-time Effects Analysis

#### A. Behavioral Engagement

*Active engagement (AE%)*. There was no significant time-on-task effect in AE% in the video watching activity (*F*(1, 58) = 0.90, *p* = 0.35, *η*^2^_p_ = 0.015). However, AE% significantly decreased in the online (*F*(1, 58) = 42.49, *p* < 0.001, *η*^2^_p_ = 0.423) and in-person activities (*F*(1, 58) = 14.34, *p* < 0.001, *η*^2^_p_ = 0.198), and significantly increased in the individual activity (*F*(1, 58) = 71.84, *p* < 0.001, *η*^2^_p_ = 0.553) (Figure 4, upper panel), suggesting engagement waned with time in the former but increased in the latter.

**Figure 4.**
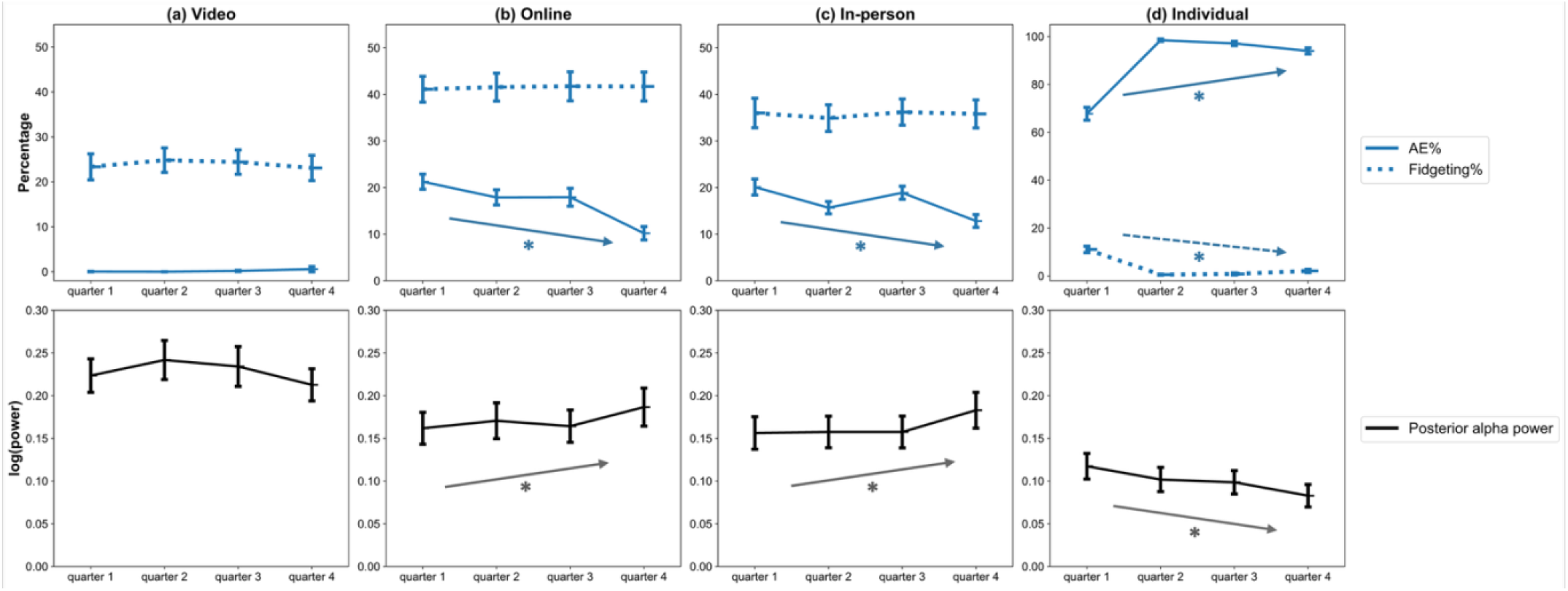
Time-on-task effects for alpha power in the posterior and central clusters (bottom row), and behavioral measures (top row), namely active engagement (AE%) and fidgeting%. During interactive activities (online, in-person) alpha power increased, and AE% decreased, whereas the opposite was observed during the student-led (individual). Notably the neural measures appeared to be related to AE% but not fidgeting, which was constant across quarters in the interactive activities, and decreases during the hands-on individual activity. No time-on-task effects were evident in the asynchronous video watching activity. *a linear contrast has significant over time (*p* < 0.05). Error bars represent standard error.

*Fidgeting*. There was no significant time-on-task effect in the video watching (*F*(1, 58) = 0.01, *p* = 0.91, *η*^2^_p_ = 0.000), online (*F*(1, 58) = 0.04, *p* = 0.84, *η*^2^_p_ = 0.001), and in-person (*F*(1, 58) = 0.00, *p* = 0.95, *η*^2^_p_ = 0.000) activities. There was a significant increase in the individual activity (*F*(1, 58) = 34.90, *p* < 0.001, *η*^2^_p_ = 0.376) (Figure 4, upper panel).

*Passive engagement (PE%)*. PE% showed no significant time-on-task effects in the video watching, online, or inperson activities, but significantly decreased during the individual activity, complementing the AE% increase.

*Off-task*. Off-task gaze/verbalizations behavior did not change significantly in the video watching, in-person, or individual activities, but increased during the online activity (c.f., supplementary materials).

#### B. EEG Features

*Alpha power*. There was no significant time-on-task effect in the video watching (*F*(1, 58) = 1.13, *p* = 0.29, *η*^2^_p_ = 0.019). However, alpha power significantly increased with time-on-task in the online (*F*(1, 58) = 3.94, *p* = 0.052, *η*^2^_p_ = 0.064) and the in-person activities (*F*(1, 58) = 3.90, *p* = 0.053, *η*^2^_p_ = 0.063), and significantly decreased in the individual activity (*F*(1, 58) = 10.70, *p* = 0.002, *η*^2^_p_ = 0.156) (Figure 4, lower panel). The results are consistent with the significant decrease in visual attention (AE%) across activity in the teacher-led (online, in-person) activities, and an increase in visual attention (AE%) over time during the individual activity.

*Spectral slope* There was no significant difference in the video watching (*F*(1, 58) = 0.42, *p* = 0.52, *η*^2^_p_ = 0.007), online (*F*(1, 58) = 1.04, *p* = 0.31, *η*^2^_p_ = 0.018), and in-person (*F*(1, 58) = 2.30, *p* = 0.14, *η*^2^_p_ = 0.038) activities over time. There was a significant flattening in the individual activity (*F*(1, 58) = 37.45, *p* < 0.001, *η*^2^_p_ = 0.392), consistent with the decrease in alpha power across quarters. See supplemental materials for plots.

In sum, time-on-task effects in neural features were absent in the video watching (online, asynchronous) activity but indicated shifts of neural state, and putatively attention, during the teacher-led (online & in-person synchronous) and student-led activities.

#### C. Within-subject Correlations

The time-on-task analysis suggested a negative relationship between neural features (alpha power, slope) and active engagement (AE%). To test this negative relationship, we fit a regression model to each pair of neural and behavioral measures within each individual and assessed the significance of the group mean regression coefficient (representing the slope) using one-sample t tests. The results of the one-sample t-tests (Figure 5(a)) showed that the regression coefficients, capturing the relationship between alpha power and AE% (*t*(58) = −7.27, *p* < 0.001, *d* = −0.95) and PE% (*t*(58) = 8.36, *p* < 0.001, *d* = 1.09), were significantly different from zero, whereby lower AE% was associated with higher alpha power (negative association), and higher PE% was also associated with higher alpha power (positive association). Complementary effects were observed for spectral slope (Figure 5(b)), with significant relationships for AE% (*t*(58) = −12.25, *p* < 0.001, *d* = −1.60), PE% (*t*(58) = 12.55, *p* < 0.001, *d* = 1.63), fidgeting (*t*(58) = 4.47, *p* < 0.001, *d* = 0.58), and off-task behavior (*t*(50) = 3.32, *p* < 0.001, *d* = 0.47). In sum, alpha power significantly tracks active engagement across time-on-task, as well as its anti-correlated counterpart, passive engagement.

**Figure 5.**
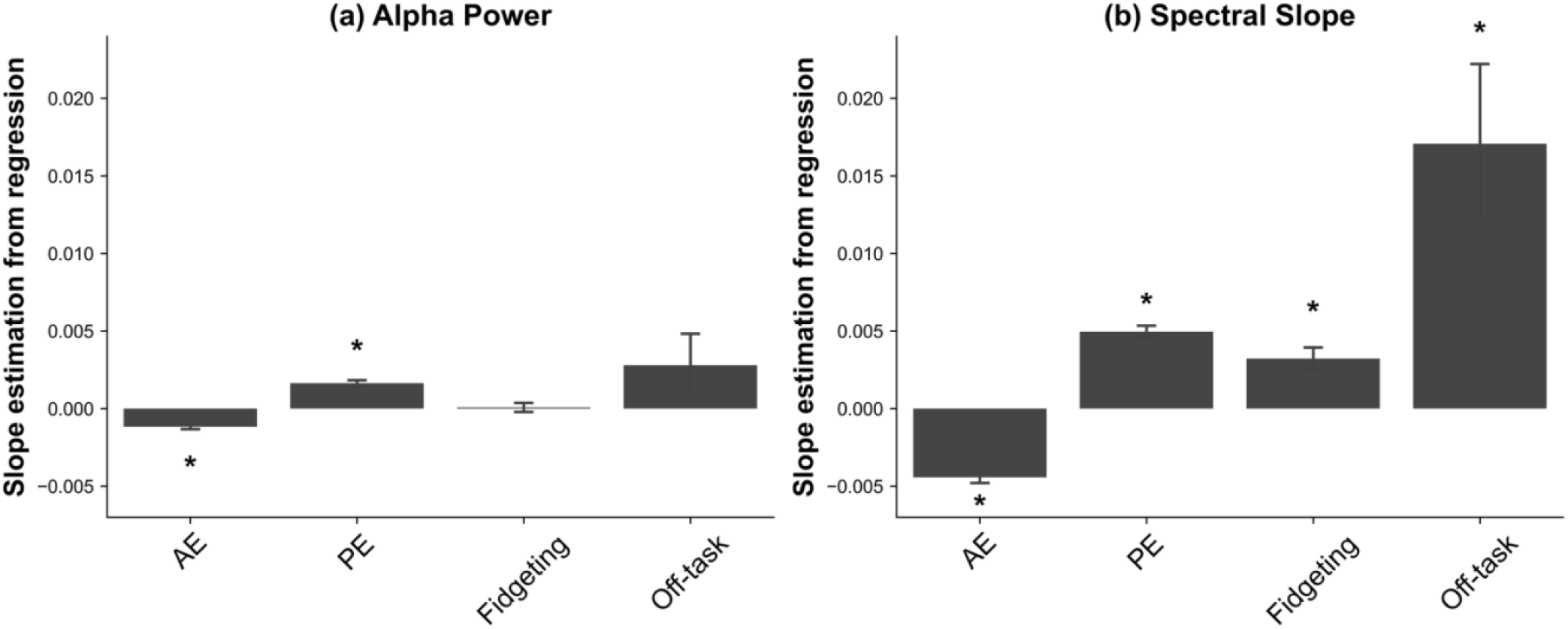
EEG-extracted feature slopes: (a) alpha power and (b) spectral slope estimated from regressions. Error bars represent standard error. * *p* < 0.001, a significant difference from zero. Error bars represent standard error.

## Discussion

In the current study, we measured behavioral and neural indices of attention while students engaged in learning activities to assess how instructional context – specifically instructional modality and management – shapes student attention, and if this differs for students with ADHD. Results revealed significant effects of both instructional modality and management on students’ engagement, across diagnostic groups. Additionally, posterior alpha power, along with spectral slope and offset, tracked active engagement (AE%) within subject. Notably, behavioral indicators of attention varied in their sensitivity to instructional context and group differences, suggesting differing mechanistic pathways.

### Instructional Context Impacts Attention Engagement

Overall, the results revealed significant effects of instructional *modality* on both neural and behavioral indices of attention engagement. Asynchronous video watching emerged as the least engaging instructional context, characterized by lower active engagement (AE%) and higher passive engagement (PE%), fidgeting, off-task behavior, as well convergently elevated alpha power. This interpretation is consistent with prior findings on video-mediated learning (Seo et al., 2021) and with reports of elevated alpha power and inattentive behaviors during video watching relative to lecture-based instruction in college students (Grammer et al., 2021). Within synchronous contexts, however, we did not observe hypothesized differences between in-person and online instruction, despite expecting such differences based on variation in instructor interaction. This pattern is consistent with prior meta-analytic work that concluded that course design and delivery are more important factors on outcome than modality (Woldeab et al., 2020). However, in the present study, the null effect could be attributed to relatively high engagement with the instructor in both online and in-person context, combined with relatively short duration of the instructional blocks (<10 min), as modality effects may emerge more clearly over longer periods. Thus, while the benefit of synchronous over asynchronous instruction is supported in the current study, the effect of online versus in-person instruction warrants further research with particular emphasis on duration and potential time-on-task interactions.

Instructional *management* also had significant effects on neural and behavioral indices of attention engagement, with student-led (individual activity) showing greater engagement relative to teacher-led (in-person activity) instruction. Counter to our prediction, this result suggests that the motivational and regulatory affordances of the hands-on, student-led activity outweighed the hypothesized benefits afforded by scaffolding provided by the instructor across diagnostic groups. This main effect was additionally supported by time-on-task effects, whereby active engagement (AE%) increased across time during student-led individual activity but decreased during both teacher-led online and in-person activities. This positive trend in student-led activity over a 10 min duration suggests that recommendations regarding attention waning in young adults after 10 minutes may be oversimplified and contingent on instruction management and task characteristics. Taken together, these findings are broadly consistent with the ISI framework (Connor et al., 2004), which posits that the extent to which attention is teacher-directed versus student-directed can change demands on children’s attentional regulation, and, thus, attention-related behavior (Connor et al., 2004).

### Individual Differences Appear in Motor Behaviors

In contrast to the robust within-subject effects of instructional context, few significant differences were observed between students with ADHD and TD peers in EEG measures during learning activities. Across instructional contexts, alpha power, spectral slope, and spectral offset (c.f., supplemental materials) did not differ reliably by group, suggesting similar patterns of visual engagement across groups during learning activities. Although this pattern may partly reflect the imbalance in sample size between the ADHD and TD groups, the absence of group differences was replicated when participants were stratified by symptom severity in more balanced subsamples (cf., supplemental materials), suggesting that activity-related demands exerted a stronger influence on measures of attention engagement than diagnostic status in our sample. Similarly, no group-by-time interactions were observed indicating similar temporal profiles across students during the activities. Prior work has reported shorter time-on-task spans in 6-11-year-old children with ADHD (2-4 min) than those without (7 min) (Rapport et al., 2009). The discrepancy could be accounted for by differences in classroom context and observational methods, both of which have been shown to moderate visual attention differences in learning settings (Kofler et al., 2008).

In contrast to the neural measures, video-coded behavioral measures, revealed group differences in a subset of behaviors. Specifically, children with ADHD exhibited higher levels of fidgeting than TD peers during synchronous teacher-led activities (online, in-person). Group differences were also observed in passive engagement (PE%), although these effects were more context-dependent and less consistent than those observed for fidgeting. In contrast, groups did not differ in active engagement (AE%) across instruction contexts, and off-task behavior (gaze shifts & verbalizations) showed a similar, but more variable, pattern of effects. Together, these findings suggest that group differences were most evident in overt motor behavior, particularly during synchronous teacher-led activities that require children to remain physically still and maintain teacher-guided attention. In complement, these data also imply that both video watching and hands-on individual activities can suppress the expression of fidgeting, though potentially by different and opposing mechanisms given that only the individual activity was associated with enhanced AE%.

### Quantifying Attentional Engagement

An unexpected but theoretically informative finding in this study was the variability in effects across neural and behavioral measures, suggesting differential sensitivity to attention related processes versus other putative sources contributing to behavior during learning. Consistent with our prior work (Grammer et al., 2021; Xu et al., 2022), alpha power, the primary neural measure examined, reliably tracked visual attention during learning activities. This interpretation is in line with prior reports of alpha power decreases tracking engagement of visual attention and associated neural substrates (Foxe & Snyder, 2011; Klimesch, 2012; Lenartowicz et al., 2025; Mathewson et al., 2011; McWhirter & Klapper, 1990). Complementing alpha power, broadly analogous effects were observed for two additional neural measures, spectral slope and offset – sometimes referred to as “aperiodic” activity. In the current study, spectral slope decreased from resting states (EC, EO) to teacher-led and student-led activities, analogous to alpha power and suggesting activity-related flattening of the power spectrum with visual engagement. This interpretation is consistent with published within-subject effects, where flatter slopes were observed with increased wakefulness or arousal (Lendner et al., 2020), with one hypothesized cortical mechanism being an increase in the ratio of excitation to inhibition (Gao et al., 2017). Notably, this interpretation differs from prior interpretations of between-subject effects of flatter spectral slope with aging (Voytek et al., 2015) and in ADHD (Pertermann et al., 2019), whereby the relative increase in higher frequencies with slope flattening was interpreted as putative neural noise. In the current study, slopes were numerically flatter in ADHD but the group difference reached significance only in the eyes closed condition (c.f., supplemental materials). In light of the within-subject slope flattening with greater engagement, this pattern could indicate that children with ADHD may be less likely to enter a rest state relative to the TD group when the eyes were closed, implying an association of spectral slope with over-activation or arousal rather than neural noise.

Finally, an important finding in this study is differential association of the video-coded behaviors to EEG features, learning contexts, and time-on-task effects, suggesting differential sensitivity to underlying mechanisms that contribute to behavioral attentiveness. In particular, whereas active engagement (AE%, i.e., following instructions) tracked effects of instructional context on neural features, this was not the case for other behavioral indicators. Namely, passive engagement (PE%), while generally negatively correlated with AE%, appeared to be elevated along with fidgeting in the video watching condition. Although fidgeting varied across activities and exhibited group-level differences, it did not covary with alpha power or AE% over time. The effects in PE% and fidgeting thus suggest that motor activity can decouple from moment-to-moment visual attention, perhaps reflecting self-regulation demands and/or individual system traits. The final metric, off-task gaze/verbalizations also did not track visual attention metrics in the current study. However, as this metric was characterized by high variance, this may be related to imprecision of coding visual gaze from video and may prove a reliable convergent measure with objective sensor data, such as from eye tracking. These data suggest a mechanistic dissociation between motor and visual behavioral indicators and highlight the value of including multiple behavioral and neural measures, as not all indices commonly assumed to reflect attention capture the same underlying processes.

### Conclusion

In sum, our findings show that instructional context—including modality and how attention is managed during learning—may play a significant role in supporting or hindering attention engagement. Furthermore, we showed that neural and behavioral measures differed in their sensitivity to visual attention versus alternate processes, with neural oscillations providing a robust index of visual engagement across contexts. The results underscore: (i) the importance of considering environmental context, such as instructional modality and management of learning, when examining mechanisms of attention and individual differences therein, and (ii) the value of recording multiple behavioral and neural measures, as not all indices commonly assumed to reflect attention capture the same underlying processes.

## Data Availability Statement

The data supporting the findings of this study are available from the corresponding author upon reasonable request, subject to institutional and ethical restrictions.

## Author Contributions

Conceptualization: JKG and AL. Data curation: FYC and AL; supporting: XZM. Formal analysis: FYC and AL; supporting: JKG. Funding acquisition: SKL, JKG, and AL. Investigation: FYC, XZM, and MK; supporting: AD. Methodology: JKG and AL; supporting: XZM and MDR. Project administration: FYC, XZM, and MK. Resources: SKL, JKG, and AL. Software: XZM. Supervision: SKL, JKG, and AL. Validation: FYC. Visualization: FYC; supporting: AL. Writing – original draft: FYC, JKG, and AL. Writing – review and editing: FYC, JKG, and AL; supporting: XZM, MK, AD, MDR, and SKL.

## Acknowledgements

We thank all participants and their caregivers. We thank Remi Torres for designing the learning activities. We thank Amna Ali, Haifa Al-Bassam, Kyle Espiritu, and Minji Kim for data collection at UCLA; Alexandra Baez, Lilly Bay, Alexa Gromada, Jordan Meyer, Sydney Saviano, and Meixin Yu for data collection and video behavioral coding at UVA; Anjali Amazigo for data collection at UVA; Chaerin Noh for EEG preprocessing support at UCLA; and Jalal Sadek for the CBCL data entry at UCLA.

## Citation Diversity Statement

Citation diversity information has been provided in accordance with the Journal of Cognitive Neuroscience guidelines.

## Declaration of and AI-assisted technologies

During the preparation of this work, the first author used ChatGPT (OpenAI) to assist with copy editing, organization of manuscript text, and troubleshooting of analysis code. The tool was not used to generate research hypotheses, interpret findings, create primary research data, or perform analyses autonomously. All analyses, methodological decisions, results, and interpretations were reviewed and verified by the authors, who take full responsibility for the published article.

