## Supplementary Materials for "Effects of Instructional Context on Neural Features of Attention during Learning Activities in Children with and without ADHD"

#### Protocol Details

##### Learning Activities Counterbalance

There were four counterbalancing sequences of learning activities designed to minimize order effects. These sequences were assigned evenly across participants, resulting in 20 participants per sequence (Table S1).

Table S1. *Counterbalancing Sequences of Learning Activities*

| **Counterbalancing Sequence** | **Activity 1** | **Activity 2** | **Activity 3** | **Activity 4** |
|---|---|---|---|---|
| Counterbalancing Sequence 1 | in-person | video | online | individual |
| Counterbalancing Sequence 2 | video | online | in-person | individual |
| Counterbalancing Sequence 3 | in-person | online | individual | video |
| Counterbalancing Sequence 4 | online | in-person | video | individual |

#### Computerized EEG Task Paradigms & Results

In addition to the learning activities, participants completed four computerized tasks: a passive auditory distractor embedded during learning activities, a laboratory-based passive auditory oddball task, a spatial working memory (SWM) task, and the Hearts and Flowers task. These tasks are not the primary focus of the present study and will be reported in future publications; they are described here for completeness. All computerized tasks were implemented using Presentation® software (Version 21, Neurobehavioral Systems, Inc., Berkeley, CA, www.neurobs.com). EEG and stimulus events were synchronized using Lab Streaming Layer (LSL; Kothe et al., 2025) via Ethernet connection (CAT6) and an Ethernet switch (NETGEAR®, model GS316).

*Passive Auditory Oddball (learning activities).* We hypothesized that distractibility would be detectable via an orienting response to background sounds. The auditory track followed an oddball structure consisting of deviant and standard events. Deviants were 2 kHz naturalistic sounds (e.g., slamming doors, knocking, honks). Standard trials were silent; thus, only the deviant sounds were audible to participants. Event distribution was randomized, resulting in deviant inter-trial intervals (ITI) ranging from 6.06–60.35 seconds (mean ITI = 20.79 seconds).

Fifty participants (ADHD:TD = 13:37) were included in analyses based on the preprocessing procedures described in the Methods section. The average number of usable epochs per activity was 25 (video), 18 (online), 15 (in-person), and 22 (individual). Due to the limited number of epochs, confirmatory analyses were not pursued. Exploratory analyses revealed no significant differences ($p > .05$) between activities when analyzed across the full sample (Figure S1(a)) or when stratified by ADHD (Figure S1(b)) and TD (Figure S1(c)) groups.

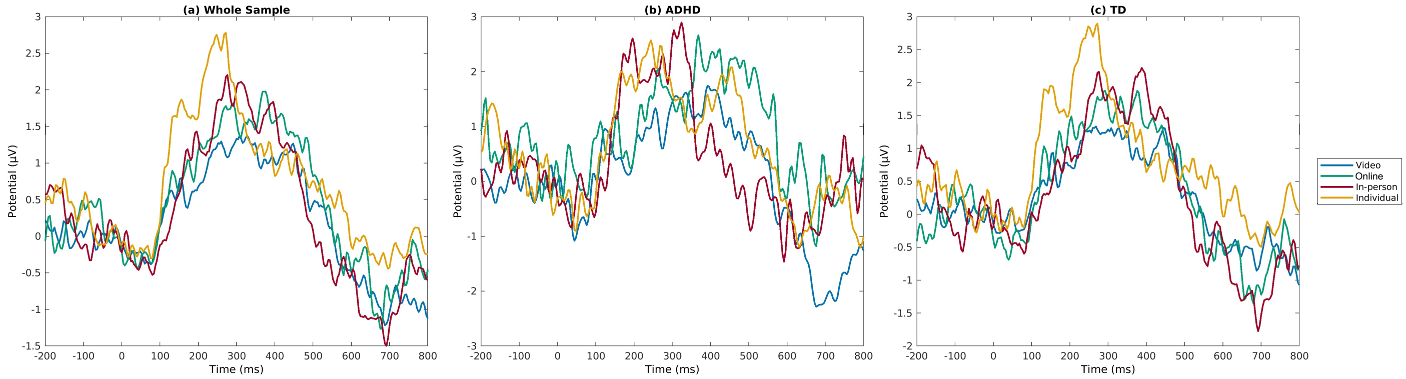

**Figure S1. ERPs during passive auditory oddball (learning activities).** (a) Whole subsample, (b) ADHD, and (c) TD. No significant differences were observed between activities ($p > 0.05$), either in the whole sample (a) or when analyses were conducted separately for the ADHD (b) and TD (c) groups. TD = typically developing.

*Passive Auditory Oddball (laboratory).* Participants viewed a child-friendly, non-stimulating video (rainfall and rainbow formation) while tones were presented. The task included 150 standard tones (1 kHz) and 50 deviant tones (2 kHz), randomized with an inter-stimulus interval (ISI) of 0.45–0.55 seconds. These data were not analyzed due to unintended event marker errors, including lack of jitter between displayed tones.

*Spatial Working Memory (SWM).* The SWM task was adapted from previous work (Lenartowicz et al., 2014) which included low-load (one yellow dot) and high-load (three yellow dots) conditions, with 40 total trials (20 per condition). Participants completed 20 trials, followed by a brief break (~1 minute), and then completed the remaining 20 trials. These results will be reported in a separate publication.

*Hearts and Flowers Task.* The Hearts and Flowers task *(Wright & Diamond, 2014)* included 12 congruent trials, 12 incongruent trials, and 33 mixed trials (congruent and incongruent). Behavioral results are presented in Table S2.

Table S2. *The Hearts and Flowers Results*

| | **ADHD** (M, SD) | **TD** (M, SD) | **Statistic** | ***p*** | **Effect size** |
|---|---|---|---|---|---|
| **Congruent (hearts)** | | | | | |
| RT (ms) | 560.53 (142.32) | 477.85 (120.05) | $t(57) = 2.20$ | 0.032 | $d = 0.66$ |
| RT sd (ms) | 150.02 (83.16) | 118.26 (75.47) | $t(57) = 1.37$ | 0.176 | $d = 0.41$ |
| Accuracy | 0.96 (0.06) | 0.96 (0.07) | $t(57) = -0.05$ | 0.961 | $d = -0.02$ |
| **Incongruent (flowers)** | | | | | |
| RT (ms) | 616.47 (159.2) | 561.5 (136.06) | $t(56) = 1.29$ | 0.203 | $d = 0.39$ |
| RT sd (ms) | 193.02 (95.16) | 129.05 (61.55) | $t(18.26) = 2.43$ | 0.026 | $d = 0.90$ |
| Accuracy | 0.89 (0.17) | 0.93 (0.11) | $t(56) = -0.98$ | 0.332 | $d = -0.29$ |
| **Congruent & incongruent mixed (hearts & flowers)** | | | | | |
| RT (ms) | 749.58 (112.16) | 741.63 (141.09) | $t(57) = 0.20$ | 0.844 | $d = 0.06$ |
| RT sd (ms) | 241.71 (124.18) | 179.12 (64.51) | $t(16.65) = 1.87$ | 0.079 | $d = 0.75$ |
| Accuracy | 0.68 (0.17) | 0.75 (0.14) | $t(57) = -1.56$ | 0.124 | $d = -0.47$ |
| **Total (hearts, flowers, mixed)** | | | | | |
| RT (ms) | 673.05 (117.82) | 636.48 (127.41) | $t(57) = 0.98$ | 0.332 | $d = 0.29$ |
| RT sd (ms) | 236.8 (79.92) | 201.71 (52.83) | $t(57) = 1.94$ | 0.058 | $d = 0.58$ |
| Accuracy | 0.79 (0.13) | 0.83 (0.1) | $t(57) = -1.32$ | 0.194 | $d = -0.39$ |

**Note.** RT = reaction time; *M* = mean; *SD* = standard deviation. Independent samples *t*-tests were used to compare group differences. Welch's *t*-tests are reported when Levene's test indicated unequal variances. Cohen's *d* is reported as the effect size. $p < 0.05$.

#### Data Processing & Feature Extraction

##### *EEG Processing*

We used custom scripts in MATLAB (The MathWorks, 2022) and EEGLAB (Delorme & Makeig, 2004) to process the data. Native .xdf EEG files were imported into EEGLAB, with electrode locations based on the default head model template (standard-10-5-cap385.elp). Data were filtered using a 0.1 Hz high-pass and 50 Hz low-pass filter. Non-stationary artifacts were attenuated using Artifact Subspace Reconstruction (ASR; Kothe, 2013) with a burst criterion of 15.

Recordings from computerized tasks and learning activities were merged prior to ICA decomposition. TP9 and TP10 electrodes were removed due to persistent muscle contamination. Remaining outlier channels were detected using flt_clean_channels.m from the ASR toolbox, based on channel correlation profiles. The initial correlation threshold was set to 0.6 and iteratively adjusted to identify between 1 and 5 bad channels per dataset. Identified bad channels were removed prior to re-referencing.

EEG data were re-referenced to the average of all remaining channels. Binary Infomax ICA (Makeig et al., 1996) was used to decompose the signal into independent components. ICLabel (Pion-Tonachini et al., 2019) was used to classify components. Components with maximal prediction confidence for muscle, eye, heart, line noise, or channel noise were removed prior to reconstruction of the cleaned time series.

Following cleaning, continuous recordings (EC, EO, and activity data) were segmented into 2-second non-overlapping epochs. Noisy epochs were identified using a 5-standard-deviation threshold and rejected using pop_autorej.m (≤3% rejection or at least one epoch). To identify dataset-level outliers, root mean square (RMS) power was calculated for each dataset across conditions. Datasets were excluded if: (i) RMS fell outside the interquartile range (IQR), (ii) more than 50% of epochs were rejected, or (iii) fewer than 10 epochs remained.

Exclusions included: age outside range (N = 1), non-participation (N = 1), missing diagnostic interview (N = 3), excessive line noise not correctable via pipeline (N = 2), >50% EEG data loss (N = 4), data structure issues due to equipment failure (N = 2), insufficient trials (<10 in baseline or activity; N = 3), and RMS outliers (N = 5). Fifty-nine participants were retained for final analyses. The average number of retained epochs was 23 (EC), 22 (EO), 242 (video), 187 (online), 169 (in-person), and 213 (individual), corresponding to an average retention rate of 82%.

##### *EEG Oscillatory Feature Extraction*

Power spectral density (PSD) was computed for each 2-second epoch using MATLAB's pwelch function without overlap. To separate oscillatory power from aperiodic components, PSD estimates were input into the spectral parameterization (specparam; formerly FOOOF; Donoghue et al., 2020) toolbox.

Specparam parameters were set as follows: peak width limits = [1, 8], maximum number of peaks = 6, minimum peak height = 0.05, and aperiodic mode = "fixed." Alpha power was defined within the 9–10 Hz subrange. This range was selected based on prior meta-analytic findings indicating alpha peaks between 9–10 Hz in pediatric samples (Freschl et al., 2022),

the central frequencies detected by specparam across resting and activity conditions (9.09–10.05 Hz), and visual inspection of the PSD (Figure S2).
Average model fit quality across selected channels was $R^2 = 0.98$ with mean error = 0.04. In addition to alpha power, spectral slope (log-transformed slope of the aperiodic component) and spectral offset were extracted.
Nine channels were selected for analysis and grouped into three clusters to improve signal-to-noise ratio while covering anterior–posterior scalp regions: frontal (F3, F4, Fz), central (C3, C4, Cz), and posterior (O1, O2, POz). Feature values were averaged within each cluster prior to statistical analysis.

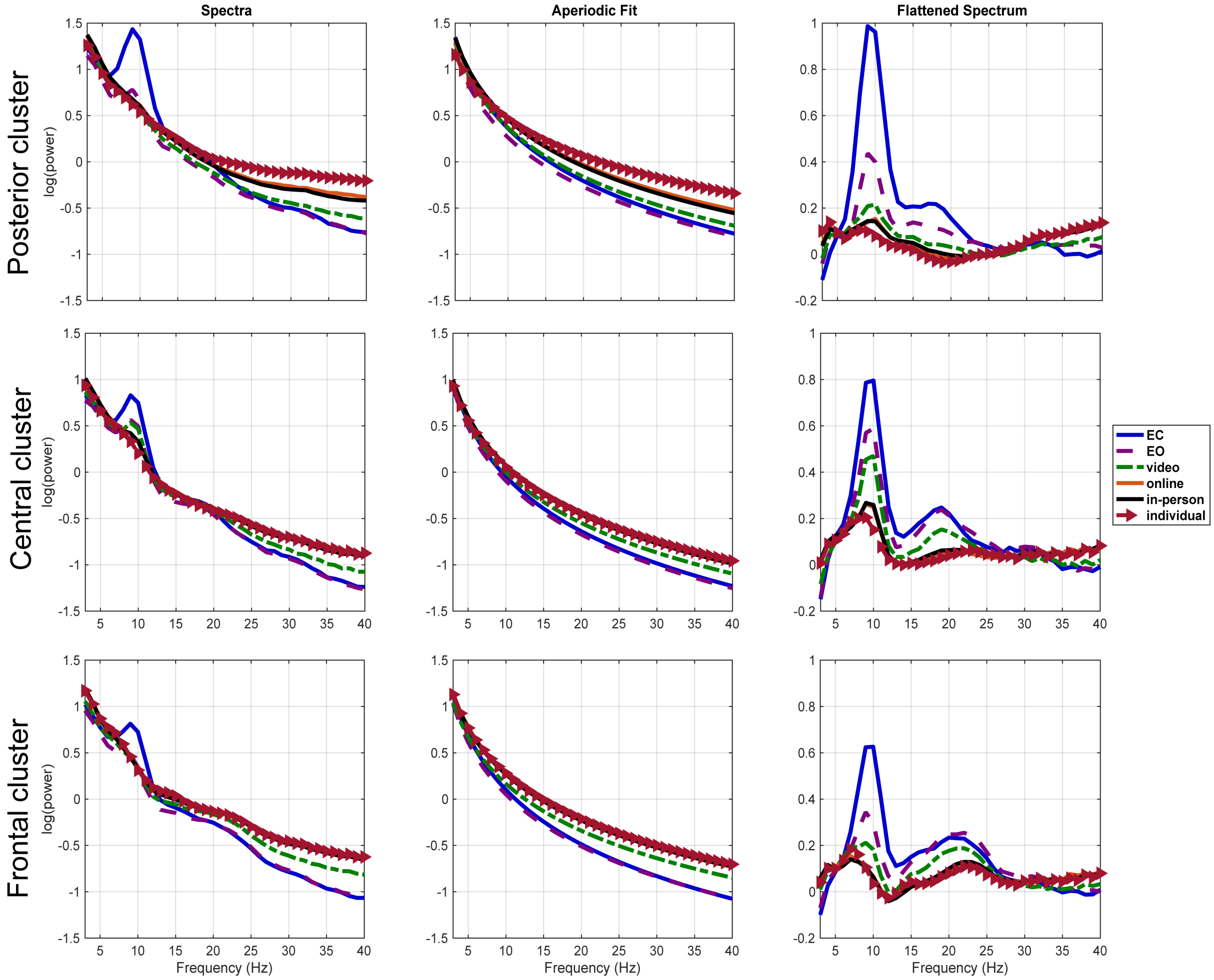

**Figure S2. The power spectrum before (a) and after (c) the specparam fitting in the subsample datasets (N=59) across 2 resting state and four activities and 3 clusters (9 channels).**

#### Behavioral Coding Method

Behavioral coding was conducted using Datavyu (Datavyu Team, 2014), an open-source video coding platform. Six undergraduate research assistants and the second author completed all coding. The Student Attention Tracking (SAT; Mao et al., under review) protocol was used, which integrates elements from previously established behavioral coding systems assessing student attention (Farley et al., 2013; Rapport et al., 2009; Shapiro, 2004). SAT included eight codes: active engagement (AE), passive engagement (PE), macro-fidgeting, micro-fidgeting, passive off-task gaze, verbal off-task verbalizations, and related subcategories (see Table 1). Raw coding data were recorded in milliseconds and subsequently converted to seconds and minutes for analysis. 25% of the sample (N = 20) was randomly selected to be double-coded by two independent coders. The overall inter-coder reliability was high (ICC = 0.905, 95% CI [0.888, 0.920]). Behavioral coding served as a convergent measure alongside EEG metrics to characterize attention-related engagement across instructional contexts.

### Clinical and Neurocognitive Assessments

##### *Neurocognitive Assessment & Diagnosis*

We assessed symptom severity using the Conners 3-Parent Short Form (Conners, 2008) and the Strengths and Weaknesses of Attention-Deficit/Hyperactivity Symptoms and Normal Behavior Scale (SWAN) (Swanson et al., 2012). Conners raw scores were converted to age- and gender-normed T-scores, and SWAN responses were scored on a −3 to 3 scale, with higher values indicating greater symptom severity. Participants with T-scores ≥ 70 on either Conners Inattention or Hyperactivity/Impulsivity subscales were referred to a licensed clinical psychologist for a semi-structured diagnostic interview with their primary caregiver. ADHD diagnoses were determined using the Schedule for Affective Disorders and Schizophrenia for School-Age Children (Kaufman et al., 1997).

To characterize broader neurocognitive and behavioral functioning, parents completed the BRIEF-2 (Gioia et al., 2015) and the CBCL (Achenbach & Rescorla, 2001), and participants completed five subtests of the Woodcock–Johnson IV (Schrank 2014a, 2014b) assessing reading, mathematics, and cognitive processing speed (Letter-Word Identification, Passage Comprehension, Applied Problems, Math Fluency, Pair Cancellation). Detailed scoring procedures and results for these measures are provided in the sections below; BRIEF-2 and CBCL findings will be examined in a future publication.

##### *Academic Achievement*

Woodcock–Johnson IV raw scores were calculated using the publisher's paper scoring form and converted to age-equivalent scores based on normative scoring tables, reflecting performance relative to the age-matched normative sample.

##### *SWAN Scoring and Results*

There were 30 items. Items 1–9 correspond to DSM ADHD Inattentive symptoms, items 10–18 to DSM Hyperactive/Impulsive symptoms, items 19–27 to Oppositional Defiant Disorder symptoms, and the final three items to symptoms commonly described as Cognitive Disengagement Syndrome (formerly Sluggish Cognitive Tempo). The results are shown in Table S3.

Table S3. *SWAN Results*

| | **ADHD** (M, SD) | **TD** (M, SD) | **Statistic** | ***p*** | **Effect size** |
|---|---|---|---|---|---|
| Inattention | 1.04 (0.63) | -0.49 (0.92) | $t(57) = 5.98$ | < 0.001 | $d = 1.79$ |
| Hyperactivity/Impulsivity | 0.93 (0.91) | -0.60 (0.79) | $t(57) = 6.19$ | < 0.001 | $d = 1.85$ |
| Oppositional Defiant Disorder | 1.01 (0.97) | -0.35 (0.73) | $t(19.76) = 4.96$ | < 0.001 | $d = 1.70$ |
| Cognitive Disengagement Syndrome | 0.40 (0.64) | -0.52 (0.80) | $t(57) = 4.06$ | < 0.001 | $d = 1.21$ |

**Note.** *M* = mean; *SD* = standard deviation. Independent samples *t*-tests were used to compare group differences. For Oppositional Defiant Disorder, Welch's *t*-test was reported because Levene's test indicated unequal variances ($p = 0.044$). Cohen's *d* is reported as the effect size. $p < 0.05$.

##### *BRIEF 2 Scoring and Results*

BRIEF 2 was scored by the publisher's online scoring platform PAR ®iConnect. Raw scores were calculated by adding up individual item responses, available for ten clinical scales, three index scores and one general score (global executive composite). There were inhibit, self-monitor, shift, emotional control, initiate, task completion, working memory, plan/organize, task-monitor, and organization of materials in the clinical scales. There were behavior regulation, emotion regulation, and cognitive regulation as three index scores. T-scores were converted from the raw scores, in relation to child's gender and age. The results are shown in Table S4.

Table S4. *BRIEF2 Results*

| | **ADHD** (M, SD) | **TD** (M, SD) | **Statistic** | ***p*** | **Effect size** |
|---|---|---|---|---|---|
| **Clinical scales** | | | | | |
| Inhibit | 67.53 (11.03) | 48.09 (8.55) | $t(57) = 7.05$ | <0.001 | $d = 2.11$ |
| Self-monitor | 61.87 (9.50) | 48.02 (8.21) | $t(57) = 5.42$ | <0.001 | $d = 1.62$ |
| Shift | 60.53 (13.49) | 51.09 (10.41) | $t(57) = 2.81$ | 0.007 | $d = 0.84$ |
| Emotional control | 64.33 (12.22) | 50.66 (9.65) | $t(57) = 4.42$ | <0.001 | $d = 1.32$ |
| Initiate | 61.40 (7.27) | 49.95 (8.69) | $t(57) = 4.58$ | <0.001 | $d = 1.37$ |
| Working memory | 68.80 (8.51) | 49.61 (8.72) | $t(57) = 7.40$ | <0.001 | $d = 2.21$ |
| Plan/Organize | 61.67 (8.80) | 49.23 (7.90) | $t(57) = 5.12$ | <0.001 | $d = 1.53$ |
| Task-monitor | 62.73 (7.06) | 49.64 (8.03) | $t(57) = 5.62$ | <0.001 | $d = 1.68$ |
| Organization of materials | 59.47 (9.90) | 50.14 (6.75) | $t(57) = 4.08$ | <0.001 | $d = 1.22$ |
| **Index Scores** | | | | | |
| Behavior regulation index (BRI) | 66.93 (9.74) | 48.02 (7.94) | $t(57) = 7.51$ | <0.001 | $d = 2.25$ |
| Emotion regulation index (ERI) | 63.60 (13.22) | 50.86 (10.02) | $t(57) = 3.91$ | <0.001 | $d = 1.17$ |
| Cognitive regulation index (CRI) | 64.47 (7.52) | 49.61 (7.57) | $t(57) = 6.57$ | <0.001 | $d = 1.97$ |
| **General score** | | | | | |
| Global executive composite (GEC) | 68.40 (9.36) | 50.05 (8.20) | $t(57) = 7.23$ | <0.001 | $d = 2.16$ |

**Note.** *M* = mean; *SD* = standard deviation. Independent samples *t*-tests were utilized to compare group differences. Cohen's *d* is reported as the effect size. $p< 0.05$.

##### *CBCL Scoring and Results*

CBCL was scored by the publisher's online scoring platform ASEBA ® (Achenbach & Rescorla, 2001). Raw scores were calculated by adding up individual item responses, available for eight syndrome scales (anxious/depressed, withdrawn/depressed, somatic complaints, social problems, thought problems, attention problems, rule-breaking behavior, aggressive behavior), three summary scores (internalizing problems, externalizing problems, total problems), six DSM-oriented scales scores (depressive problems, anxiety problems, somatic problems, attention deficit, oppositional defiant problems, conduct problems), three competence scale scores (activities, social, school), and three 2007 scale scores (sluggish cognitive tempo, obsessive-compulsive problems, stress problems). T scores were converted from the raw scores, in relation to child's gender and age. The results are shown in Table S5.

Table S5. *CBCL Results*

| | **ADHD** (M, SD) | **TD** (M, SD) | **Statistic** | ***p*** | **Effect size** |
|---|---|---|---|---|---|
| **Syndrome scales** | | | | | |
| Anxious/Depressed | 62.73 (10.23) | 58.35 (7.94) | $t(57) = 1.72$ | 0.091 | $d = 0.52$ |
| Withdrawn/Depressed | 60.00 (7.17) | 56.12 (6.46) | $t(57) = 1.91$ | 0.062 | $d = 0.57$ |
| Somatic complaints | 59.73 (8.73) | 55.23 (5.96) | $t(18.55) = 1.84$ | 0.082 | $d = 0.67$ |
| Social problems | 59.60 (7.14) | 53.58 (3.89) | $t(16.86) = 3.10$ | 0.007 | $d = 1.23$ |
| Thought problems | 63.13 (9.46) | 54.14 (4.86) | $t(16.57) = 3.50$ | 0.003 | $d = 1.42$ |
| Attention problems | 69.13 (7.62) | 54.12 (4.70) | $t(57) = 8.52$ | <0.001 | $d = 2.55$ |
| Rule-breaking behavior | 62.33 (7.47) | 53.30 (4.48) | $t(17.53) = 4.38$ | <0.001 | $d = 1.67$ |
| Aggressive behavior | 66.60 (11.28) | 52.74 (4.70) | $t(15.74) = 4.56$ | <0.001 | $d = 1.97$ |
| **Summary scores** | | | | | |
| Internalizing problems | 61.40 (11.81) | 55.95 (9.01) | $t(57) = 1.84$ | 0.070 | $d = 0.55$ |
| Externalizing problems | 63.07 (13.66) | 48.42 (8.34) | $t(17.66) = 3.87$ | 0.001 | $d = 1.47$ |
| Total problems | 64.93 (11.89) | 51.35 (7.70) | $t(18.16) = 4.09$ | <0.001 | $d = 1.51$ |
| **DSM-oriented scales** | | | | | |
| Depressive problems | 61.53 (7.87) | 55.19 (6.47) | $t(57) = 3.08$ | 0.003 | $d = 0.92$ |
| Anxiety problems | 64.67 (10.41) | 58.33 (8.20) | $t(57) = 2.32$ | 0.024 | $d = 0.69$ |
| Somatic problems | 57.93 (10.27) | 55.05 (6.31) | $t(17.68) = 0.99$ | 0.334 | $d = 0.38$ |
| Attention deficit | 69.27 (7.09) | 52.74 (4.05) | $t(18.08) = 8.29$ | <0.001 | $d = 3.07$ |
| Oppositional defiant problems | 64.93 (8.46) | 53.49 (4.56) | $t(17.56) = 4.79$ | <0.001 | $d = 1.82$ |
| Conduct problems | 64.40 (8.66) | 53.14 (5.61) | $t(18.52) = 4.58$ | <0.001 | $d = 1.66$ |

**Note.** *M* = mean; *SD* = standard deviation. Independent-samples *t*-tests were utilized to compare group differences. When Levene's test indicated unequal variances ($p < .05$), Welch's *t*-test was reported. Cohen's *d* is reported as the effect size. $p< 0.05$.

#### Statistical Analyses

##### *Group Statistics*

Repeated-measures ANOVAs were conducted with activity (four levels) as a within-subject factor and diagnostic group as a between-subject factor. Gender was initially included as a covariate in all models. Because no significant activity × gender interactions were observed across EEG features or behavioral measures, gender was removed from the remaining analyses. Given the unequal sample sizes between participants with ADHD (N = 15) and typically developing participants (TD; N = 44), a secondary grouping approach was implemented using symptom severity thresholds. Participants were classified into high- and low-symptom groups based on T-scores ≥ 60 on inattentive and hyperactivity subscales, resulting in balanced groups (high: N = 29; low: N = 30). Identical repeated-measures ANOVA models (activity × symptom group) were applied to this alternative grouping to assess the stability of effects under balanced sample conditions.

##### *Brain-Behavior Analyses*

To evaluate time-on-task effects, each learning activity was divided into four equal temporal quarters. For each quarter, alpha power, spectral slope, and spectral offset were extracted from the predefined channel clusters. Specparam model fitting was conducted separately for each quarter and channel. Across the nine channels and four quarters, the average model fit was $R^2 = 0.97$ with a mean fitting error of 0.04. Behavioral coding percentages were calculated using the same quarter-based segmentation procedure. Post hoc linear contrasts were used to evaluate linear trends across quarters within each activity.

To assess within-subject brain–behavior associations, individual linear regressions were computed for each participant by regressing video-coded behavioral outcomes on alpha power across the entire session (collapsed across all four activities). The resulting regression coefficients were retained as subject-level slope estimates. These slopes were then evaluated at the group level using one-sample t-tests to determine whether the mean slope differed from zero.

Finally, additional analyses were conducted to evaluate potential confounding factors. Chi-square tests were used to assess group differences in gender distribution. Independent-samples t-tests were conducted to examine group differences in questionnaire subscales (Conners, SWAN, WJ), activity duration, and the number of usable EEG epochs. The significance threshold was set at $\alpha = .05$ for all analyses. Statistical analyses were performed using IBM SPSS Statistics (IBM Corp., 2022).

#### EEG Cluster Results

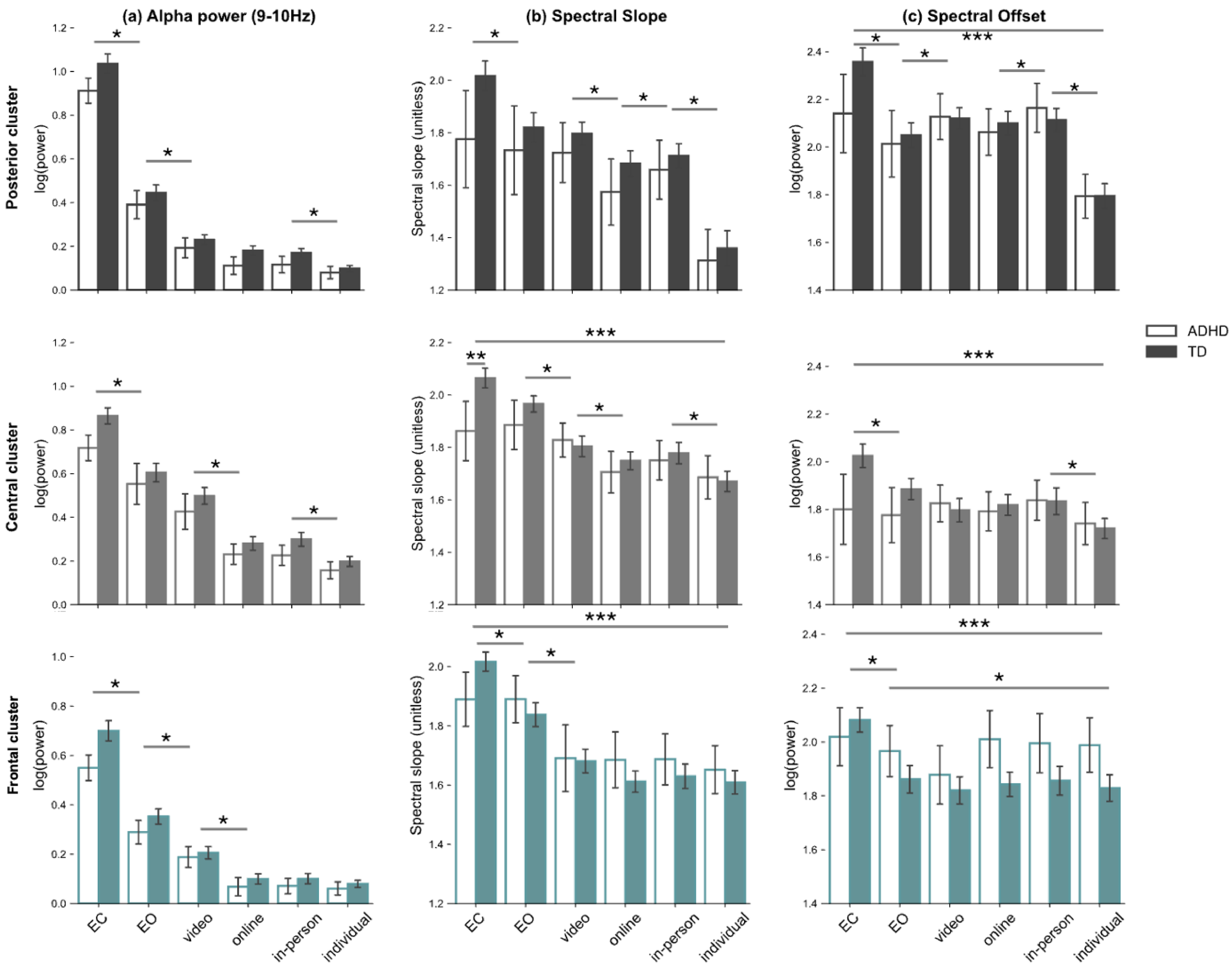

**Figure S3. Average (a) alpha power, (b) spectral slope, and (c) spectral offset across posterior (O1, O2, POz; top row), central (C3, C4, Cz; middle row), and frontal (F3, F4, Fz; bottom row) electrode clusters.** Across analyses, main effects of group were not significant unless otherwise noted. Alpha power (a) decreased from the resting state (EC > EO), to asynchronous (EO > video watching), to teacher-led synchronous (video watching > online/in-person), and finally to student-led (individual) activities. A similar pattern was observed for spectral slope (b), suggesting a flattening of the power spectrum accompanying the decrease in alpha power. Effects were more variable in the spectral offset parameter (c). In the central cluster, slope and offset showed an interaction driven by elevated values in the TD group during EC (see main text for details). * $p < 0.05$, main effect of activity. ** $p < 0.05$, main effect of group. *** $p < 0.05$, interaction between activity and group. TD = typically developing. Error bars represent standard error.

#### Alpha Power

*Central Cluster.* The central cluster showed effects similar to those observed in the posterior cluster. Neither the main effect of group ($F(1,55) = 1.99$, $p = 0.163$, $\eta^2_p = 0.035$), nor the interaction between activity and group was significant ($F(5,275) = 0.57$, $p = 0.72$, $\eta^2_p = 0.010$). The central cluster showed only a significant main effect of activity ($F(5,275) = 3.18$, $p = 0.008$, $\eta^2_p = 0.055$), as alpha power decreased across activities (from left to right in the middle panel of Figure S3(a)). The post-hoc pairwise results indicated that alpha power was the highest in EC relative to EO and learning activities ($p < 0.001$). Alpha power was also higher in EO relative to all learning activities ($p < 0.001$). Among instructional contexts, alpha power was highest during the video watching activity ($p < 0.001$) relative to other activities, and lowest during the individual activity ($p < 0.001$) relative to other activities. Again, the post-hoc pairwise results showed that the alpha power in the online and in-person conditions was not significantly different ($p > 0.05$).

*Frontal Cluster.* The frontal cluster showed effects analogous to the posterior and central clusters, except that no main effect of activity was observed ($F(5,275) = 0.91$, $p = 0.47$, $\eta^2_p = 0.016$). Neither the main effect of group ($F(1,55) = 1.03$, $p = 0.31$, $\eta^2_p = 0.018$) nor the interaction between activity and group ($F(5,275) = 0.85$, $p = 0.51$, $\eta^2_p = 0.015$) was significant. Alpha power decreased across two resting-state conditions and learning activities (from left to right in the lower panel of Figure S3(a)). Post-hoc pairwise results indicated that alpha power was the highest in EC relative to all other activities ($p < 0.001$). Alpha power was also higher in EO relative to all learning activities ($p < 0.001$). Among the activities, alpha power was the highest during the video watching activity ($p < 0.001$) relative to other activities and the lowest during the individual activity ($p < 0.001$) relative to other activities. However, the post-hoc pairwise results showed that alpha power between online and in-person or between in-person and individual activities was not significantly different ($p > 0.05$).

#### Spectral Slope

*Central Cluster.* In the central cluster, the spectral slope was analogous to that observed in the posterior cluster, and there was no main effect of group ($F(1,55) = 1.01$, $p = 0.31$, $\eta^2_p = 0.018$). There was a significant main effect of activity ($F(5,275) = 2.53$, $p = 0.029$, $\eta^2_p = 0.044$). The slope decreased from the least to most engaging activities (from left to right in the middle panel of Figure S3(b)). The slope was also higher in EO relative to learning activities ($p < 0.05$). Among activities, the slope was highest during the video watching activity ($p < 0.001$) relative to other activities, except for the in-person activity ($p = 0.085$), and the lowest during the individual activity ($p < 0.001$), except for the online activity ($p = 0.098$). However, the post-hoc pairwise results showed that the slope between the online and in-person was not significantly different ($p = 0.177$). Finally, and unlike for the posterior cluster, the interaction between activity and group was significant ($F(5,275) = 2.85$, $p = 0.020$, $\eta^2_p = 0.049$). A post-hoc independent samples t-test showed that the slope was significantly higher in the TD group ($M = 2.06$, $SD = 0.25$) than in the ADHD group ($M = 1.86$, $SD = 0.44$), $t(57) = -2.21$, $p = 0.032$, Cohen's $d = 0.56$ in EC, but not in EO and the four learning activities ($p > 0.05$).

*Frontal Cluster.* In the frontal cluster, there was no main effect of group ($F(1,55) = 0.20$, $p = 0.65$, $\eta^2_p = 0.004$). There was a significant interaction between activity and group ($F(5,275) = 2.27$, $p = 0.048$, $\eta^2_p = 0.040$). The slope decreased from the least to most engaging activities ($F(5,275) = 4.21$, $p = 0.001$, $\eta^2_p = 0.071$) (from left to right in the lower panel of Figure S3(b)). Post-hoc pairwise results indicated that the slope was the highest (slope was steepest) in EC relative to all other activities ($p < 0.05$). The slope was also higher in EO relative to all learning activities ($p < 0.001$). However, there was no significant difference in the slope among the four learning activities ($p > 0.05$).

#### Spectral Offset

*Posterior Cluster.* Finally, we evaluated spectral offset (indicative of broadband power), which showed qualitatively analogous, albeit statistically less reliable, effects. There was no main effect of group ($F(1,55) = 0.76$, $p = 0.38$, $\eta^2_p = 0.014$) or activity ($F(5,275) = 2.02$, $p = 0.075$, $\eta^2_p = 0.036$). However, the interaction between activity and group was significant ($F(5,275) = 2.32$, $p = 0.044$, $\eta^2_p = 0.040$), as shown in the upper panel of Figure S3(c). Post-hoc pairwise comparisons indicated that offset was highest during EC relative to all activities ($p < 0.05$), except for video watching and in-person activities ($ps > 0.05$). Offset was also higher during the in-person activity than during the online and individual activities ($ps < 0.05$), but did not differ from the remaining conditions ($p > 0.05$). Finally, offset was lowest during individual activity relative to all other activities ($p < 0.001$).

*Central Cluster.* Similar to the posterior cluster, there was no main effect of group ($F(1,55) = 1.09$, $p = 0.30$, $\eta^2_p = 0.019$). There was a significant interaction between activity and group ($F(5,275) = 3.40$, $p = 0.005$, $\eta^2_p = 0.058$). Unlike the posterior cluster, there was a trend-level effect of activity ($F(5,275) = 2.20$, $p = 0.055$, $\eta^2_p = 0.038$), as shown in the middle panel of Figure S3(c). Post-hoc pairwise comparisons indicated that offset was highest during EC relative to all activities ($p < 0.05$), but did not significantly differ from the in-person activity ($p > 0.05$). Offset was second highest during the in-person activity; however, it did not significantly differ from EO or the other activities ($p > 0.05$), except for the individual activity ($p = 0.01$). Among the learning activities, offset was highest during in-person teaching, but this difference was not statistically significant ($p > 0.05$), except relative to the individual activity ($p = 0.01$). Finally, offset was lowest during the individual activity ($p < 0.05$).

*Frontal Cluster.* Similar to the posterior and central clusters, there was no main effect of group ($F(1,55) = 0.70$, $p = 0.403$, $\eta^2_p = 0.013$). There was a significant interaction between activity and group ($F(5,275) = 2.66$, $p = 0.023$, $\eta^2_p = 0.046$), consistent with both the posterior and central clusters. Unlike the posterior cluster, but similar to the central cluster, the frontal cluster showed a significant main effect of activity ($F(5,275) = 2.72$, $p = 0.020$, $\eta^2_p = 0.047$), as shown in the lower panel of Figure S3(c). Post-hoc pairwise comparisons indicated that offset was highest during EC relative to all other activities ($p < 0.05$). In contrast to the posterior and central clusters, no other pairwise comparisons were significant ($p > 0.05$).

#### Age Effects in EEG Features

There were between-subject effects of age on posterior spectral slope ($F(1,55) = 5.30$, $p = 0.025$, $\eta^2_p = 0.088$), posterior offset ($F(1,55) = 6.98$, $p = 0.011$, $\eta^2_p = 0.113$), and central offset ($F(1,55) = 5.69$, $p = 0.021$, $\eta^2_p = 0.094$). Older participants showed flatter spectral slope and lower offset. In addition, there was a within-subject age * activity interaction for frontal alpha power ($F(5,275) = 7.05$, $p < 0.001$, $\eta^2_p = 0.114$), frontal slope ($F(5,275) = 3.04$, $p = 0.01$, $\eta^2_p = 0.052$), and frontal offset ($F(5,275) = 2.50$, $p = 0.031$, $\eta^2_p = 0.043$).

### Task-on-Time Effects Analysis

#### Behavioral Engagement

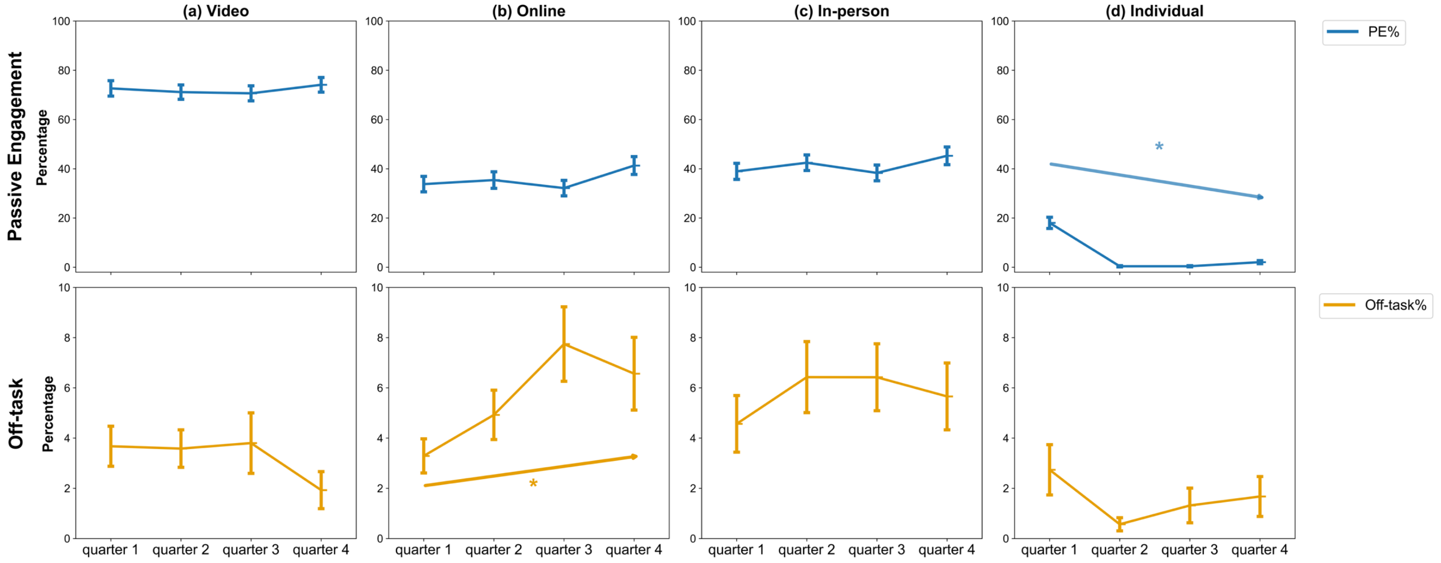

**Figure S4. Time-on-task effects for behavioral measures across learning activities.** Behavioral measures include passive engagement (PE%; upper row) and off-task behavior (lower row). During the student-led (individual) activity, PE% decreased significantly over time. Significant time-on-task effects were observed for off-task behavior during the online activity, which increased over time. * indicates a significant linear contrast over time ($p < .05$). Error bars represent standard error.

*Passive engagement (PE%).* There was no significant time-on-task effect in the video watching ($F(1, 58) = 0.14$, $p = 0.71$, $\eta^2_p = 0.002$), online ($F(1, 58) = 3.42$, $p = 0.07$, $\eta^2_p = 0.056$), and in-person ($F(1, 58) = 1.60$, $p = 0.21$, $\eta^2_p = 0.027$) activities. There was a significant decrease time-on-task in the individual activity ($F(1, 58) = 48.55$, $p < 0.001$, $\eta^2_p = 0.456$), complementing the increase in AE% (Figure S4, upper row).

*Off-task.* There was no significant time-on-task effect in the video watching ($F(1, 58) = 2.66$, $p = 0.11$, $\eta^2_p = 0.044$), in-person ($F(1, 58) = 1.45$, $p = 0.23$, $\eta^2_p = 0.024$), and individual ($F(1, 58) = 0.66$, $p = 0.42$, $\eta^2_p = 0.011$) activities. There was a significant increase in the online activity ($F(1, 58) = 6.80$, $p = 0.012$, $\eta^2_p = 0.105$) (Figure S4, lower row).

#### EEG Extracted Features

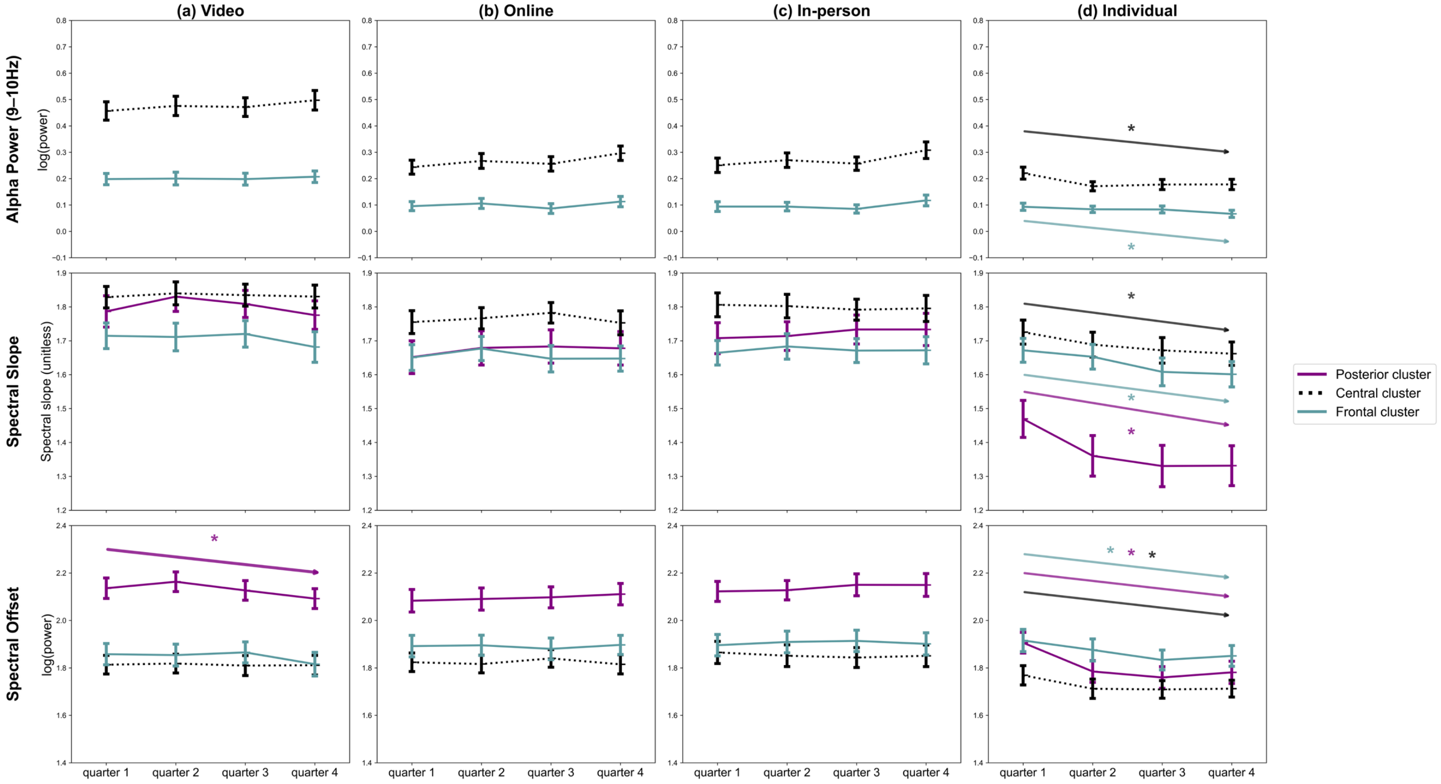

**Figure S5. Time-on-task effects for alpha power (top row), spectral slope (middle row), and spectral offset (bottom row) in the frontal cluster.** Alpha power (9–10 Hz) is shown for the central and frontal clusters. Spectral slope and spectral offset are shown for the posterior (purple), central (black dotted), and frontal (teal) clusters. During the student-led (individual) activity, alpha power in the frontal cluster decreased significantly over time. Significant time-on-task effects were also observed during the student-led activity for spectral slope and spectral offset across clusters, with slope and offset decreasing over time. In addition, posterior spectral offset during the video activity showed a significant time-on-task effect. No other significant time-on-task effects were observed during the video, online, or in-person activities. * indicates a significant linear contrast over time ($p < .05$). Error bars represent standard error.

*Alpha Power.* Across clusters, there was no significant time-on-task effect in the video watching activity ($F(1, 58)_{central} = 2.10$, $p = 0.15$, $\eta^2_p = 0.035$; $F(1, 58)_{frontal} = 0.30$, $p = 0.58$, $\eta^2_p = 0.005$). However, alpha power significantly increased with time-on-task in the online ($F(1, 58)_{central} = 8.92$, $p = 0.004$, $\eta^2_p = 0.133$) and in-person activities ($F(1, 58)_{central} = 5.80$, $p = 0.02$, $\eta^2_p = 0.091$), but not in the frontal cluster (online: $F(1, 58) = 0.99$, $p = 0.32$, $\eta^2_p = 0.017$; on-person: $F(1, 58) = 1.50$, $p = 0.23$, $\eta^2_p = 0.025$). In addition, alpha power significantly decreased in the individual activity ($F(1, 58)_{central} = 10.76$, $p = 0.002$, $\eta^2_p = 0.157$; $F(1, 58)_{frontal} = 10.00$, $p = 0.002$, $\eta^2_p = 0.147$) (Figure S5, first row).

*Spectral Slope.* Across clusters, there was no significant time-on-task effect in the video watching ($F(1, 58)_{central} = 0.000$, $p = 1.000$, $\eta^2_p = 0.000$; $F(1, 58)_{frontal} = 1.22$, $p = 0.27$, $\eta^2_p = 0.021$), online ($F(1, 58)_{central} = 0.01$, $p = 0.91$, $\eta^2_p = 0.000$; $F(1, 58)_{frontal} = 0.22$, $p = 0.64$, $\eta^2_p = 0.004$), or in-person ($F(1, 58)_{central} = 0.41$, $p = 0.53$, $\eta^2_p = 0.007$; $F(1, 58)_{frontal} = 0.02$, $p = 0.90$, $\eta^2_p = 0.000$) activities. There was a significant flattening of the spectral slope in the individual activity ($F(1, 58)_{central} = 9.25$, $p = 0.004$, $\eta^2_p = 0.138$; $F(1, 58)_{frontal} = 12.27$, $p < 0.001$, $\eta^2_p = 0.175$), consistent with the decrease in alpha power across quarters (Figure S5, second row).

*Spectral Offset.* Across clusters, there was no significant time-on-task effect in the online ($F(1, 58)_{posterior} = 1.06$, $p = 0.31$, $\eta^2_p = 0.018$; $F(1, 58)_{central} = 0.000$, $p = 0.979$, $\eta^2_p = 0.000$; $F(1, 58)_{frontal} = 0.00$, $p = 0.997$, $\eta^2_p = 0.000$) and in-person ($F(1, 58)_{posterior} = 2.53$, $p = 0.12$, $\eta^2_p = 0.042$; $F(1, 58)_{central} = 0.38$, $p = 0.54$, $\eta^2_p = 0.007$; $F(1, 58)_{frontal} = 0.06$, $p = 0.81$, $\eta^2_p = 0.001$) activities. There was a significant decrease during video watching only in the posterior cluster ($F(1, 58)_{posterior} = 4.06$, $p = 0.049$, $\eta^2_p = 0.065$), and in the other clusters, the video watching effect was not significant ($F(1, 58)_{central} = 0.06$, $p = 0.81$, $\eta^2_p = 0.001$; $F(1, 58)_{frontal} = 3.12$, $p = 0.08$, $\eta^2_p = 0.051$). There was a significant decrease in the individual activity across clusters ($F(1, 58)_{posterior} = 26.92$, $p < 0.001$, $\eta^2_p = 0.317$; $F(1, 58)_{central} = 6.63$, $p = 0.013$, $\eta^2_p = 0.103$; $F(1, 58)_{frontal} = 10.49$, $p = 0.002$, $\eta^2_p = 0.153$) (Figure S5, third row).

### Within-Subject Regression Coefficients Statistical Results

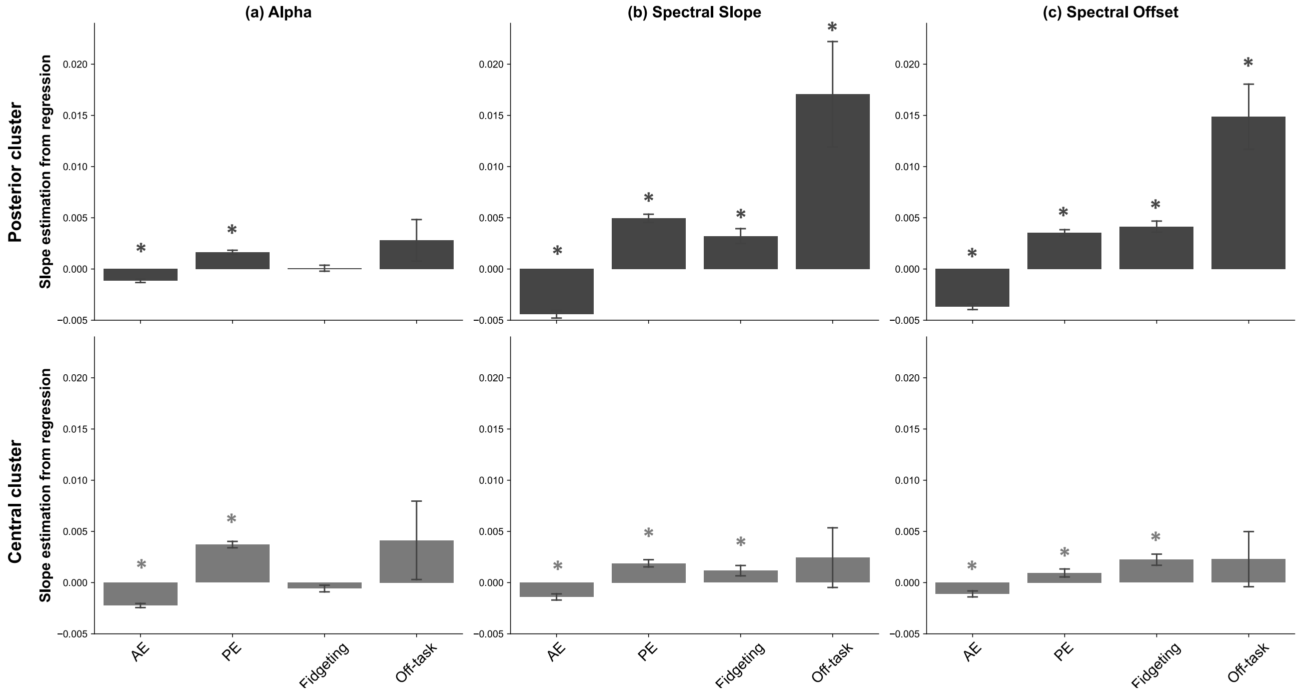

**Figure 6. EEG-extracted feature slopes: (a) alpha, (b) spectral slope, (c) spectral offset estimated from regressions.** Error bars represent standard error. * *p* < 0.001, a significant difference from zero. Error bars represent standard error.

To further characterize within-subject brain–behavior associations, linear regression models were fit within each individual, pairing neural measures (alpha power, spectral slope, and spectral offset) with behavioral indices (AE%, PE%, fidgeting, and off-task). The resulting regression coefficients were tested against zero using one-sample t-tests. Consistent with the main text, in the central cluster, alpha power was negatively associated with AE% and positively associated with PE% (p < .001), with no significant associations for fidgeting or off-task behavior. In analyses of spectral slope and offset, there were significant differences for AE%, PE%, and fidgeting in the central cluster (p < .001). However, a significant difference for off-task behavior was not observed in the central cluster (p > .40). For the posterior cluster, spectral offset showed significant associations with AE%, PE%, and fidgeting (p < .001). Table S6 presents the mean and standard deviation of the estimated slope values for posterior and central alpha power.

Table S6. *Results of One-Sample Tests of Within-Subject Regression Coefficients*

| | **Slope value** (M, SD) | **Statistic** | ***p*** | **Effect size** |
|---|---|---|---|---|
| **Posterior alpha power** | | | | |
| Active engagement (AE) | -0.001160 (0.001226) | *t*(58) = -7.27 | <0.001 | *d* = -0.95 |
| Passive engagement (PE) | 0.001634 (0.001502) | *t*(58) = 8.36 | <0.001 | *d* = 1.09 |
| Fidgeting | 0.000078 (0.002237) | *t*(58) = 0.27 | 0.791 | *d* = 0.04 |
| Off-task | 0.002795 (0.014521) | *t*(50) = 1.38 | 0.175 | *d* = 0.19 |
| **Central alpha power** | | | | |
| Active engagement (AE) | -0.002235 (0.001540) | *t*(58) = -11.16 | <0.001 | *d* = -1.45 |
| Passive engagement (PE) | 0.003713 (0.002382) | *t*(58) = 11.97 | <0.001 | *d* = 1.56 |
| Fidgeting | -0.000578 (0.002494) | *t*(58) = -1.78 | 0.080 | *d* = -0.23 |
| Off-task | 0.004131 (0.027340) | *t*(50) = 1.08 | 0.286 | *d* = 0.15 |

**Note.** *M* = mean; *SD* = standard deviation. A one-sample *t*-test was used to examine whether the average estimated slope differed significantly from zero. Cohen's *d* is reported as the effect size. *p* < .05.

### Symptom Group Analyses

This was in part motivated by unequal sample sizes in the diagnosis-based groups. The re-grouping was based on Conners scores: participants with either a Conners Inattention or Hyperactivity T score ≥ 60 were assigned to the high symptom group, and the remaining participants were assigned to the low symptom group. To further validate the grouping, correlation analyses were performed to examine the relationship between Conners and SWAN subscales. Results showed that the Conners Inattention scale was significantly correlated with the SWAN Inattention scale ($r$ = 0.701, $p$ < 0.001), and the Conners Hyperactivity scale was significantly correlated with the SWAN Hyperactivity scale ($r$ = 0.695, $p$ < 0.001). All diagnosis-based analyses reported in the main text were repeated using symptom group in place of diagnostic category. The results largely replicated the diagnostic-label effects reported in the main text and are summarized below. Tables S6–S10 present descriptive statistics and group comparisons for demographic characteristics, questionnaire measures, and task performance.

#### Participants, Neurocognitive Assessment & Diagnosis, Academic Achievement

There were 29 participants in the high symptom group and 31 in the low symptom group in the final analysis (Table S7). There were no significant differences in sex or age between the two groups ($p$ = 0.795 and $p$ = 0.808, respectively). Regarding ADHD ratings, the Conners Inattention and Hyperactivity subscales were significantly higher in the high symptom group than in the low symptom group ($p$ < .001). There were no significant differences in Woodcock–Johnson scores ($p$ > 0.05).

**Descriptive Results**

Table 2. *Participant Demographics*

| | **High symptom** (M, SD) | **Low symptom** (M, SD) | **Statistic** | ***p*** | **Effect size** |
|---|---|---|---|---|---|
| N | 29 | 30 | - | - | - |
| Sex (male : female) | 17:12 | 16:14 | $\chi^2(1) = 0.17$ | 0.683 | $V = 0.05$ |
| Age (year) | 8.57 (1.36) | 8.48 (1.39) | $t(57) = 0.25$ | 0.808 | $d = 0.06$ |
| **ADHD Rating Scale Measure** | | | | | |
| Conners - Inattention scale | 72.86 (13.57) | 48.33 (6.04) | $t(38.37) = 8.92$ | < 0.001 | $d = 2.35$ |
| Conners - Hyperactivity scale | 69.14 (12.82) | 47.53 (5.54) | $t(37.85) = 8.35$ | < 0.001 | $d = 2.20$ |
| **Academic achievement (WJ)** | | | | | |
| Letter-Word Identification | 50.41 (14.82) | 51.63 (16.90) | $t(57) = -0.29$ | 0.770 | $d = -0.08$ |
| Passage Comprehension | 27.97 (8.25) | 27.47 (9.53) | $t(57) = 0.22$ | 0.831 | $d = 0.06$ |
| Applied Problems | 29.69 (7.10) | 31.00 (6.16) | $t(57) = -0.82$ | 0.418 | $d = -0.21$ |
| Math Facts Fluency | 43.45 (29.11) | 47.10 (20.77) | $t(57) = -0.56$ | 0.580 | $d = -0.15$ |
| Pair Cancellation | 42.65 (15.74) | 42.43 (12.57) | $t(57) = 0.06$ | 0.952 | $d = 0.02$ |

**Note.** N = number; M = mean; SD = standard deviation. $\chi^2$ tests were used to examine sex differences between groups; independent-samples t tests were used to examine group differences in age, ADHD symptom measures, and WJ scores. Effect sizes are reported as Cramér's $V$ for $\chi^2$ tests and Cohen's $d$ for t tests.

Table S3. *SWAN Results*

| | **High symptom** (M, SD) | **Low symptom** (M, SD) | **Statistic** | ***p*** | **Effect size** |
|---|---|---|---|---|---|
| Inattention | 0.44 (0.98) | -0.63 (0.90) | $t(57) = 4.39$ | < 0.001 | $d = 1.14$ |
| Hyperactivity/Impulsivity | 0.42 (0.93) | -0.81 (0.79) | $t(57) = 5.51$ | < 0.001 | $d = 1.44$ |
| Oppositional Defiant Disorder | 0.43 (1.07) | -0.43 (0.70) | $t(57) = 3.65$ | < 0.001 | $d = 0.95$ |
| Cognitive Disengagement Syndrome | 0.01 (0.80) | -0.58 (0.82) | $t(57) = 2.79$ | 0.007 | $d = 0.73$ |

**Note.** $M$ = mean; $SD$ = standard deviation. Independent-samples $t$-tests were used to compare group differences. Cohen's $d$ is reported as the effect size. $p < .05$.

Table S9. *BRIEF2 Results*

| | **High symptom** (M, SD) | **Low symptom** (M, SD) | **Statistic** | ***p*** | **Effect size** |
|---|---|---|---|---|---|
| **Clinical scales** | | | | | |
| Inhibit | 61.83 (11.55) | 44.53 (5.66) | $t(40.40) = 7.26$ | <0.001 | $d = 1.91$ |
| Self-monitor | 56.59 (11.04) | 46.67 (7.06) | $t(57) = 4.13$ | <0.001 | $d = 1.08$ |
| Shift | 58.83 (12.38) | 48.33 (8.89) | $t(57) = 3.75$ | <0.001 | $d = 0.98$ |
| Emotional control | 60.45 (12.84) | 48.03 (6.69) | $t(41.83) = 4.63$ | <0.001 | $d = 1.22$ |
| Initiate | 58.00 (9.66) | 47.90 (6.79) | $t(50.10) = 4.63$ | <0.001 | $d = 1.21$ |
| Working memory | 62.62 (10.96) | 46.63 (6.61) | $t(45.72) = 6.76$ | <0.001 | $d = 1.77$ |
| Plan/Organize | 58.10 (9.40) | 46.87 (6.33) | $t(57) = 5.40$ | <0.001 | $d = 1.41$ |
| Task-monitor | 58.07 (9.30) | 48.03 (7.14) | $t(57) = 4.66$ | <0.001 | $d = 1.21$ |
| Organization of materials | 56.52 (8.57) | 48.63 (6.78) | $t(57) = 3.93$ | <0.001 | $d = 1.02$ |
| **Index Scores** | | | | | |
| Behavior regulation index (BRI) | 60.83 (11.25) | 45.1 (5.41) | $t(39.98) = 6.80$ | <0.001 | $d = 1.79$ |
| Emotion regulation index (ERI) | 60.41 (12.88) | 48.0 (7.58) | $t(45.04) = 4.49$ | <0.001 | $d = 1.18$ |
| Cognitive regulation index (CRI) | 59.93 (9.45) | 47.07 (5.20) | $t(43.20) = 6.45$ | <0.001 | $d = 1.69$ |
| **General score** | | | | | |
| Global executive composite (GEC) | 62.76 (11.03) | 46.93 (5.19) | $t(39.53) = 7.01$ | <0.001 | $d = 1.85$ |

**Note.** *M* = mean; *SD* = standard deviation. Independent-samples *t*-tests were utilized to compare group differences. When Levene's test indicated unequal variances ($p < .05$), Welch's *t*-test was reported. Cohen's *d* is reported as the effect size. $p< .05$.

Table S10. *CBCL Results*

| | **High symptom** (M, SD) | **Low symptom** (M, SD) | **Statistic** | ***p*** | **Effect size** |
|---|---|---|---|---|---|
| **Syndrome scales** | | | | | |
| Anxious/Depressed | 60.71 (8.51) | 58.33 (8.88) | $t(57) = 1.03$ | 0.306 | $d = 0.27$ |
| Withdrawn/Depressed | 58.29 (6.81) | 56.03 (6.73) | $t(57) = 1.34$ | 0.185 | $d = 0.35$ |
| Somatic complaints | 57.79 (8.06) | 55.10 (5.65) | $t(50.51) = 1.48$ | 0.145 | $d = 0.39$ |
| Social problems | 56.89 (6.43) | 53.50 (4.02) | $t(47.22) = 2.40$ | 0.020 | $d = 0.63$ |
| Thought problems | 59.57 (8.67) | 53.57 (4.52) | $t(42.27) = 3.34$ | 0.002 | $d = 0.88$ |
| Attention problems | 63.93 (8.52) | 52.47 (3.65) | $t(37.94) = 6.83$ | <0.001 | $d = 1.80$ |
| Rule-breaking behavior | 58.75 (7.75) | 52.73 (3.65) | $t(39.90) = 3.81$ | <0.001 | $d = 1.00$ |
| Aggressive behavior | 61.04 (10.77) | 51.93 (4.22) | $t(36.44) = 4.30$ | <0.001 | $d = 1.13$ |
| **Summary scores** | | | | | |
| Internalizing problems | 59.43 (9.56) | 55.43 (10.16) | $t(57) = 1.58$ | 0.121 | $d = 0.41$ |
| Externalizing problems | 57.61 (13.16) | 47.17 (7.59) | $t(44.96) = 3.73$ | <0.001 | $d = 0.98$ |
| Total problems | 60.14 (10.9) | 49.93 (7.88) | $t(51.38) = 4.14$ | <0.001 | $d = 1.09$ |
| **DSM-oriented scales** | | | | | |
| Depressive problems | 58.39 (7.66) | 55.37 (6.84) | $t(57) = 1.63$ | 0.109 | $d = 0.42$ |
| Anxiety problems | 61.00 (9.16) | 59.00 (9.22) | $t(57) = 0.94$ | 0.351 | $d = 0.25$ |
| Somatic problems | 56.96 (8.99) | 54.70 (5.83) | $t(48.25) = 1.19$ | 0.238 | $d = 0.31$ |
| Attention deficit | 63.39 (8.83) | 51.07 (2.03) | $t(30.95) = 7.51$ | <0.001 | $d = 1.99$ |
| Oppositional defiant problems | 60.61 (8.54) | 52.57 (3.85) | $t(38.66) = 4.81$ | <0.001 | $d = 1.27$ |
| Conduct problems | 59.07 (9.25) | 53.23 (5.81) | $t(47.17) = 3.02$ | 0.004 | $d = 0.79$ |

**Note.** *M* = mean; *SD* = standard deviation. Independent-samples *t*-tests were utilized to compare group differences. When Levene's test indicated unequal variances ($p < .05$), Welch's *t*-test was reported. Cohen's *d* is reported as the effect size. $p< .05$.

Table S11. *The Hearts and Flowers Results*

| | **High symptom** (M, SD) | **Low symptom** (M, SD) | **Statistic** | ***p*** | **Effect size** |
|---|---|---|---|---|---|
| **Congruent (hearts)** | | | | | |
| RT (ms) | 487.07 (133.42) | 510.28 (127.76) | $t(57) = -0.68$ | 0.498 | $d = -0.18$ |
| RT sd (ms) | 115.78 (76.07) | 136.54 (79.79) | $t(57) = -1.02$ | 0.311 | $d = -0.27$ |
| Accuracy | 0.97 (0.05) | 0.96 (0.08) | $t(57) = 0.72$ | 0.473 | $d = 0.19$ |
| **Incongruent (flowers)** | | | | | |
| RT (ms) | 566.36 (142.64) | 585.07 (145.31) | $t(56) = -0.50$ | 0.623 | $d = -0.13$ |
| RT sd (ms) | 153.51 (86.22) | 137.68 (65.36) | $t(56) = 0.79$ | 0.435 | $d = 0.21$ |
| Accuracy | 0.93 (0.12) | 0.92 (0.13) | $t(56) = 0.17$ | 0.867 | $d = 0.04$ |
| **Congruent & incongruent mixed (hearts & flowers)** | | | | | |
| RT (ms) | 719.17 (101.44) | 767.32 (156.52) | $t(49.92) = -1.41$ | 0.166 | $d = -0.36$ |
| RT sd (ms) | 208.04 (99.60) | 182.45 (72.22) | $t(57) = 1.13$ | 0.262 | $d = 0.30$ |
| Accuracy | 0.75 (0.16) | 0.72 (0.14) | $t(57) = 0.72$ | 0.474 | $d = 0.19$ |
| **Total (hearts, flowers, mixed)** | | | | | |
| RT (ms) | 628.89 (105.46) | 662.11 (141.37) | $t(57) = -1.02$ | 0.312 | $d = -0.27$ |
| RT sd (ms) | 214.73 (66.91) | 206.67 (57.8) | $t(57) = 0.50$ | 0.622 | $d = 0.13$ |
| Accuracy | 0.83 (0.11) | 0.81 (0.11) | $t(57) = 0.92$ | 0.362 | $d = 0.24$ |

**Note.** *M* = mean; *SD* = standard deviation. Independent-samples *t*-tests were utilized to compare group differences. When Levene's test indicated unequal variances ($p < .05$), Welch's *t*-test was reported. Cohen's *d* is reported as the effect size. $p< .05$.

#### EEG Extracted Features

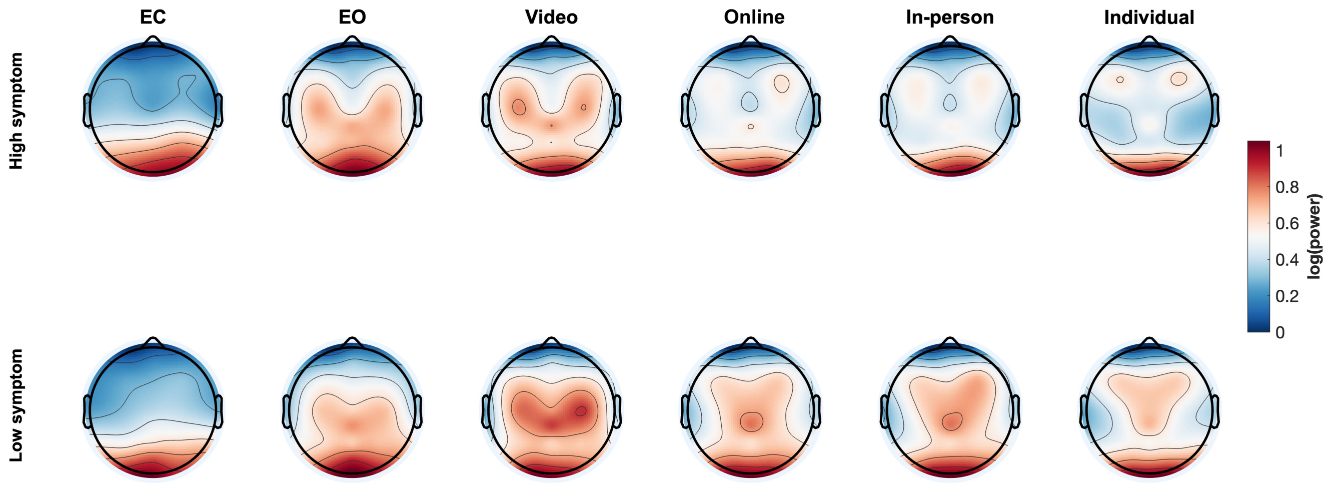

**Figure S7. Topographical distribution of alpha power across resting state and activities divided in groups.** Alpha was elevated during the resting state (EC, EO) relative to activities, with an occipital scalp distribution. During learning activities, alpha power was observed across both central and posterior scalp regions, decreasing progressively from asynchronous (video watching) to teacher-led synchronous (online, in-person) to student-led (individual) activities.

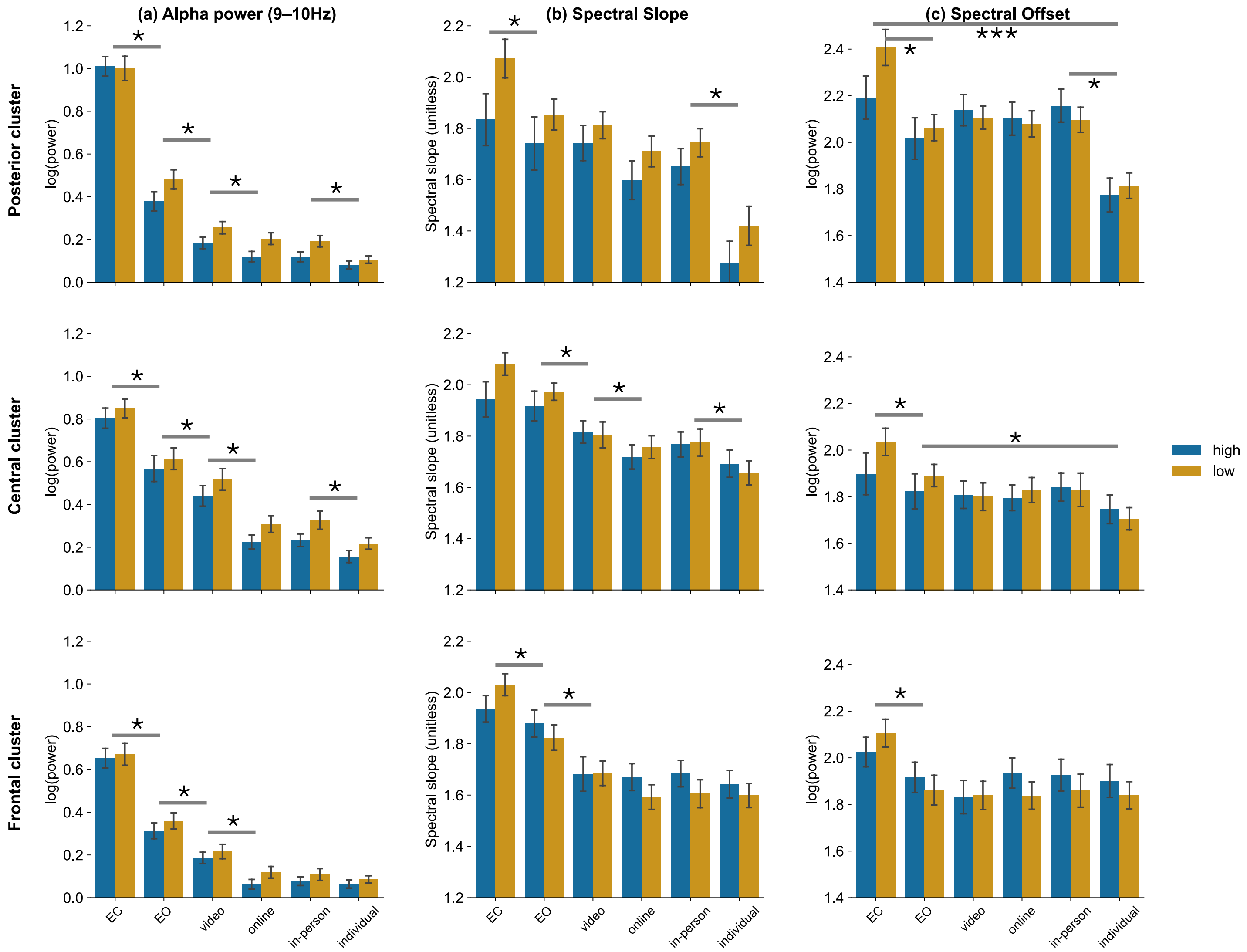

**Figure S8. Average (a) alpha power, (b) slope, and (c) offset in posterior (O1, O2, POz; upper row), central (C3, C4, Cz; middle row), and frontal (F3, F4, Fz; bottom row) electrode clusters.** Across analyses, main effects of group were not significant. Alpha power (left column) decreased from the resting state (EC > EO), to asynchronous (EO > video watching), to teacher-led synchronous (video watching > online / in-person), to student-led (individual) activities. A similar pattern was observed for the slope (middle column), suggesting a flattening of the power spectrum with alpha power decrease. Effects were more variable in the offset parameter (right column). Additionally, in the posterior cluster, the offset showed an interaction, resulting from elevated values in the low symptom group during the EC (see main text for details). ** $p < 0.05$, main effect of activity. ** $p < 0.05$, main effect of group. *** $p < 0.05$, interaction between activity and group. Error bars represent standard error.

#### Alpha Power

*Posterior Cluster.* In the posterior cluster, neither the main effect of group ($F(1,57) = 2.63$, $p = 0.11$, $\eta^2_p = 0.044$) nor the interaction between activity and group was significant ($F(5,285) = 1.31$, $p = 0.26$, $\eta^2_p = 0.022$). The posterior cluster showed only a significant main effect of activity ($F(5,285) = 344.02$, $p < 0.001$, $\eta^2_p = 0.858$), as alpha power decreased across the two resting-state conditions and learning activities (from left to right in the upper panel of Figure S8(a)). Post-hoc pairwise results indicated that alpha power was the highest in EC relative to all learning activities ($p < 0.001$). Alpha power was also higher in EO relative to all activities ($p < 0.001$). Among the activities, alpha power was the highest during the video watching activity ($p < 0.001$) relative to other activities, and lowest during the individual activity ($p < 0.001$) relative to other activities. However, the post-hoc pairwise results showed that alpha power did not significantly differ between the online and in-person activities ($p = 1.00$).

*Central Cluster*. The central cluster showed effects analogous to the posterior cluster. Neither the main effect of group ($F(1,57) = 1.31$, $p = 0.26$, $\eta^2_p = 0.022$) nor the interaction between activity and group was significant ($F(5,285) = 0.45$, $p = 0.89$, $\eta^2_p = 0.006$). The central cluster also showed only a significant main effect of activity ($F(5,285) = 180.71$, $p < 0.001$, $\eta^2_p = 0.760$), as alpha power decreased across the two resting-state conditions and learning activities (from left to right in the middle panel of Figure S8(a)). Post-hoc pairwise results indicated that alpha power was the highest in EC relative to all other activities ($p < 0.001$). Alpha power was also higher in EO relative to all learning activities ($p < 0.001$). Among the activities, alpha power was the highest during the video watching activity ($p < 0.001$) relative to other activities, and the lowest during the individual activity ($p < 0.001$) relative to other activities. However, the post-hoc pairwise results showed that alpha power did not significantly differ between the online and in-person activities ($p = 1.00$).

*Frontal Cluster*. The frontal cluster showed effects analogous to the posterior and central clusters. Neither the main effect of group ($F(1,57) = 0.58$, $p = 0.45$, $\eta^2_p = 0.010$) nor the interaction between activity and group was significant ($F(5,285) = 0.23$, $p = 0.95$, $\eta^2_p = 0.004$). The frontal cluster showed only a significant main effect of activity ($F(5,285) = 182.77$, $p < 0.001$, $\eta^2_p = 0.762$), as alpha power decreased across the two resting-state conditions and learning activities (from left to right in the bottom panel of Figure S8(a)). Post-hoc pairwise results indicated that alpha power was the highest in EC relative to all other activities ($p < 0.001$). Alpha power was also higher in EO relative to all learning activities ($p < 0.001$). Among the activities, alpha power was highest during the video watching activity ($p < 0.001$) relative to the other learning activities. However, alpha power did not significantly differ among the online, in-person, and individual activities (all *ps* = 1.00).

#### Spectral Slope

*Posterior Cluster***.** The spectral slope, which represents the log slope of the spectrum, showed results consistent with those of alpha power. In the posterior cluster, neither the main effect of group ($F(1,57) = 1.87$, $p = 0.18$, $\eta^2_p = 0.032$) nor the interaction between activity and group was significant ($F(5,285) = 1.24$, $p = 0.29$, $\eta^2_p = 0.021$). As with alpha power, the posterior cluster showed only a significant main effect of activity, with the slope decreasing across the least to most engaging conditions ($F(5,285) = 58.13$, $p < 0.001$, $\eta^2_p = 0.505$) (from left to right in the upper panel of Figure S8(b)), which is consistent with a flattening of the spectral slope. Post-hoc pairwise results indicated that the slope was the steepest in EC relative to all other activities ($p < 0.005$). The slope was also higher in EO relative to all learning activities ($p < 0.05$), except for video watching and in-person ($p > 0.05$) activities. Among activities, the slope was the highest (steepest) during the video watching activity ($p < 0.01$) relative to other activities and the lowest during the individual activity ($p < 0.001$) relative to other activities. However, post-hoc pairwise results showed that the slope between online and in-person was not significantly different ($p = 0.41$).

*Central Cluster*. In the central cluster, the spectral slope was analogous to that in the posterior cluster. Neither the main effect of group ($F(1,57) = 0.30$, $p = 0.58$, $\eta^2_p = 0.005$) nor the interaction between activity and group was significant ($F(5,285) = 2.00$, $p = 0.08$, $\eta^2_p = 0.034$). There was a significant main effect of activity ($F(5,285) = 35.01$, $p < 0.001$, $\eta^2_p = 0.381$). The slope decreased from the least to most engaging conditions (from left to right in the middle panel of Figure S8(b)). Post-hoc pairwise results indicated that the slope was the highest (slope was steepest) in EC relative to all other activities ($p < 0.001$), except for EO ($p = 0.14$). Again, the slope was also higher in EO relative to the learning activities ($p < 0.001$). Among the learning activities, the slope was highest during the video watching activity, differing significantly from the online ($p = 0.043$) and individual ($p < 0.001$) activities, but not from the in-person activity ($p = 1.00$). There were no significant differences among the online, in-person, and individual activities (*ps* > 0.05).

*Frontal Cluster*. In the frontal cluster, neither the main effect of group ($F(1,57) = 0.20$, $p = 0.66$, $\eta^2_p = 0.003$) nor the interaction between activity and group was significant ($F(5,285) = 2.02$, $p = 0.08$, $\eta^2_p = 0.034$). The slope decreased from the least to most engaging conditions ($F(5,285) = 40.42$, $p < 0.001$, $\eta^2_p = 0.415$) (from left to right in Figure S8(c)). Post-hoc pairwise results indicated that the slope was steepest in EC relative to all other activities ($p < 0.001$). The slope was also higher in EO relative to all learning activities ($p < 0.001$). However, there was no significant difference in the slope among the four learning activities ($p > 0.05$).

#### Spectral Offset

*Posterior Cluster*. In the posterior cluster, there was no main effect of group ($F(1,57) = 0.17$, $p = 0.68$, $\eta^2_p = 0.003$). However, there was a main effect of activity ($F(5,285) = 32.34$, $p < 0.001$, $\eta^2_p = 0.362$), as shown in the top panel of Figure S8(c), and an interaction between activity and group ($F(5,285) = 3.15$, $p = 0.009$, $\eta^2_p = 0.052$). Post-hoc pairwise results indicated that offset was the highest in the EC relative to all other conditions ($p < 0.05$), except for in-person ($p = 0.06$). Offset was significantly higher in the in-person activity than in the individual activity ($p < 0.001$), but did not significantly differ from the other activities ($p > 0.05$). Offset in EO was significantly higher than in the individual activity ($p < 0.001$), but did not significantly differ from the other activities ($p > 0.05$). Although the activity × group interaction was significant, follow-up comparisons revealed only a trend toward lower offset in the high-symptom group during EC (two-sided $p = 0.070$), and no simple effects reached significance.

*Central Cluster*. In the central cluster, neither the main effect of group ($F(1,57) = 0.16$, $p = 0.69$, $\eta^2_p = 0.003$) nor the interaction between activity and group was significant ($F(5,285) = 1.60$, $p = 0.16$, $\eta^2_p = 0.027$). Similar to the posterior cluster, the central cluster showed only a significant main effect of activity ($F(5,285) = 9.52$, $p < 0.001$, $\eta^2_p = 0.143$), as shown in the middle panel of Figure S8(c). Post-hoc pairwise results indicated that offset was the highest in the EC relative to all other activities ($p < 0.05$), except for the in-person activity ($p = 0.20$). Offset was the second highest in the EO, significantly greater than in the individual activity ($p = 0.003$), but not significantly different from the other activities ($p > 0.05$). Among the learning activities, offset was highest during the in-person activity and lowest during the individual activity; however, these differences were not statistically significant ($p > 0.05$).

*Frontal Cluster*. In the frontal cluster, neither the main effect of group ($F(1,57) = 0.16$, $p = 0.69$, $\eta^2_p = 0.003$) nor the interaction between activity and group was significant ($F(5,285) = 1.88$, $p = 0.098$, $\eta^2_p = 0.032$). The frontal cluster showed only a significant main effect of activity ($F(5,285) = 11.65$, $p < 0.001$, $\eta^2_p = 0.17$), as shown in the bottom panel of Figure S8(c). Post-hoc pairwise results indicated that offset was highest in EC relative to all other activities ($p < 0.001$).

#### Behavioral Engagement

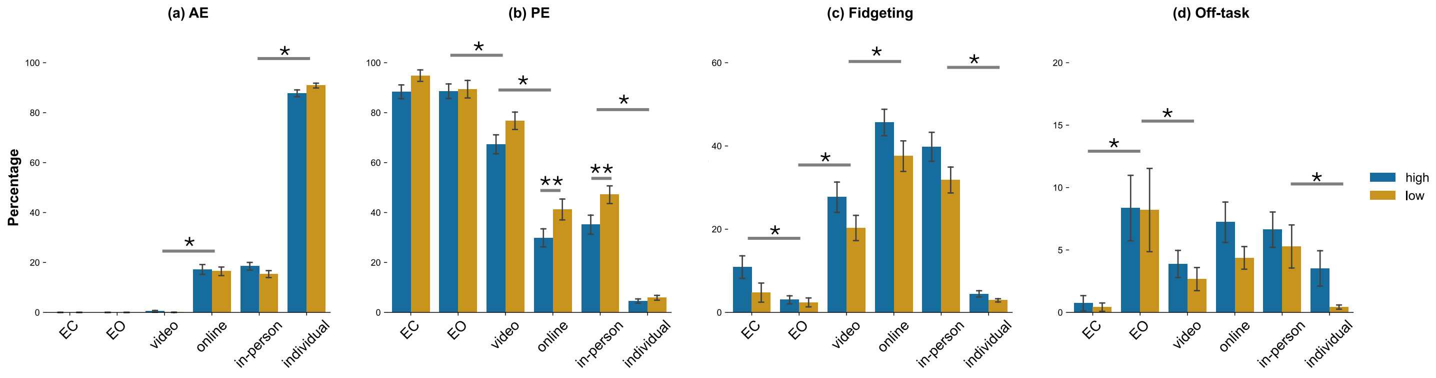

**Figure S9. Video-coded behavior across instruction context.** Percentage of each coded behavior category in each activity and group. Main effects of activity are consistent with neural measures, with the highest level of active engagement (AE) and the lowest levels of passive engagement (PE), fidgeting, and off-task gaze/vocalizations observed during the self-guided (individual) activity. * $p < 0.05$, main effect of activity. ** $p < 0.05$, main effect of group. Error bars represent standard error.

*Active Engagement (AE%)*. Neither the main effect of group ($F(1,57) = 0.08$, $p = 0.78$, $\eta^2_p = 0.001$) nor the interaction between activity and group was significant ($F(5,285) = 1.78$, $p = 0.12$, $\eta^2_p = 0.030$). However, consistent with spectral EEG features, there was a significant main effect of activity ($F(5,285) = 2163.62$, $p < 0.001$, $\eta^2_p = 0.974$). AE% increased across the two resting-state conditions and learning activities (from left to right in Figure S9(a)). Post-hoc pairwise results indicated that AE% was the highest in the individual activity relative to all other activities ($p < 0.001$). AE% was also higher in the teacher-led learning activities (online, in-person) relative to video watching, EO, and EC ($p < 0.001$), and there was no significant difference between AE% in the online and in-person activities ($p > 0.05$). The results are consistent with the EEG data, suggesting that attention engagement is the strongest in the student-led activity, followed by teacher-led activities (online, in-person), and finally asynchronous online learning (video watching).

*Passive Engagement (PE%).* Consistent with AE%, there was a main effect of activity ($F(5,285) = 329.82$, $p < 0.001$, $\eta^2_p = 0.853$); PE% decreased across the two resting-state conditions and learning activities (left to right in the upper panel of Figure S9(b)). There was no interaction between group and activity ($F(5,285) = 1.71$, $p = 0.13$, $\eta^2_p = 0.028$). Post-hoc pairwise results showed that PE% was highest in EC relative to the other activities ($p < 0.001$), but did not significantly differ from EO ($p = 1.00$). PE% was also higher in EO relative to the learning activities ($p < 0.001$). Regarding PE% in activities, video watching had the highest PE% among all activities ($p < 0.001$), while the individual activity had the lowest PE% ($p < 0.001$). Unlike AE%, there was a significant group effect ($F(1,57) = 5.84$, $p = 0.02$, $\eta^2_p = 0.093$) for PE%. As shown in Figure S9(b), post-hoc t-tests showed that the high symptom group had lower PE% than the low symptom group in the online ($p = 0.04$) and in-person ($p = 0.03$) activities. Thus, PE% showed an overall trend opposite to AE% across activities and, unlike AE%, differentiated groups in teacher-led (online, in-person) activities.

*Fidgeting.* There was no significant interaction between activity and group ($F(5,285) = 1.02$, $p = 0.41$, $\eta^2_p = 0.018$) in fidgeting measures. However, there was a significant main effect of group ($F(1,57) = 6.03$, $p = 0.017$, $\eta^2_p = 0.096$) and activity ($F(5,285) = 104.68$, $p < 0.001$, $\eta^2_p = 0.647$) (Figure S9(c)). Post-hoc pairwise results showed that fidgeting was the highest in the online activity relative to other activities ($p < 0.001$), except for in-person activity ($p = 0.11$). Fidgeting during the in-person activity was also elevated relative to other activities ($p < 0.001$), except for online activity ($p = 0.11$). There was no significant difference in fidgeting between the EC, EO, and individual activities ($p > 0.05$). In sum, fidgeting behavior showed effects analogous to, but opposite in direction from, the PE% metric, suggesting greater fidgeting (and lower PE%) during teacher-led, synchronous activities (online, in-person) and asynchronous learning (video watching) than during the student-led (individual) activity or resting-state conditions (EC, EO).

*Off-Task Gaze/Vocalizations.* For measures of off-task gaze and vocalizations, neither the main effect of group ($F(1,57) = 1.16$, $p = 0.29$, $\eta^2_p = 0.020$) nor the interaction between activity and group was significant ($F(5,285) = 0.25$, $p = 0.94$, $\eta^2_p = 0.004$). However, there was a significant main effect of activity ($F(5,285) = 8.01$, $p < 0.001$, $\eta^2_p = 0.123$) (Figure S9(d)). Post-hoc pairwise results showed that EC had the lowest level of off-task behaviors relative to all other activities ($p < 0.05$), except for the individual activity ($p > 0.05$). Off-task behaviors in the individual activity were significantly lower relative to the in-person learning activity ($p = 0.002$). There was no significant difference between online and in-person activities ($p > 0.05$), and off-task behaviors were not significantly different between video watching and online activities ($p = 0.09$).
